# Translatome and translation dynamics analysis of a RiboCancer cell line panel reveals that leukemia-associated Rps15 mutations rewire translation through codon-specific tRNA accommodation defects

**DOI:** 10.1101/2025.11.18.687986

**Authors:** Anaïs Astier, Marino Caruso, Stijn Vereecke, Paulo E. Santo, Coralie Capron, Carine Froment, Dana Rinaldi, David Cabrerizo Granados, Marine Leclercq, Jonathan Royaert, Jelle Verbeeck, Naomy Pasau, Laura Plassart, Daniele Pepe, Steven Verbruggen, Hermes Paraqindes, Simon Lebaron, Virginie Marcel, Sébastien Durand, Clément Chapat, Gerben Menschaert, Pierre Close, Francesca Rapino, Frédéric Catez, Julien Marcoux, Célia Plisson-Chastang, Kim De Keersmaecker

## Abstract

Deletions and point mutations targeting ribosomal proteins (RPs) have been identified in cancer. Yet, their role in translational dysregulation remains poorly understood. We performed an integrated genome-wide translatome analysis (proteome, Ribo-seq and total RNA-seq) as well as RiboMethSeq on an isogenic cell line library modeling the most recurrent RP defects in cancer (Rpl5^+/−^, Rpl11^+/−^, Rpl22^+/−^, Rpl22^−/−^, Rpl10 R98S, Rps15 P131S and Rps15 H137Y). RP knock-out had minimal effects on translation, whereas RP point mutations induced a significant number of translation efficiency changes, affecting up to 10% of expressed genes in Rps15 mutants associated with Chronic Lymphocytic Leukemia (CLL). Cryo-electron microscopy and biochemical analyses revealed that the Rps15 mutations destabilize the C-terminal Rps15 domain, affecting the translation elongation cycle dynamics, and deregulating accommodation of aminoacylated tRNAs at the ribosomal A-site. Using Ribo-seq and translation reporter assays, we show that this accommodation defect shows codon specificity, explaining the reduced translation efficiency of genes enriched for these codons in Rps15 mutant cells, such as histones. Notably, genes with reduced translation efficiency in Rps15 mutated cells were enriched for transcriptional regulators such as transcription factor Runx3, resulting in downregulation of Runx3 target genes involved in immune regulation. Altogether, this study provides a comparative map of the translational rewiring driven by the most frequent somatic RP mutations. We provide unprecedented mechanistic insights in the translation defects induced by CLL-associated Rps15 mutations, and reveal an unappreciated cross-talk between translational and transcriptional dysregulation in these RP mutant cells.

**GRAPHICAL ABSTRACT:** 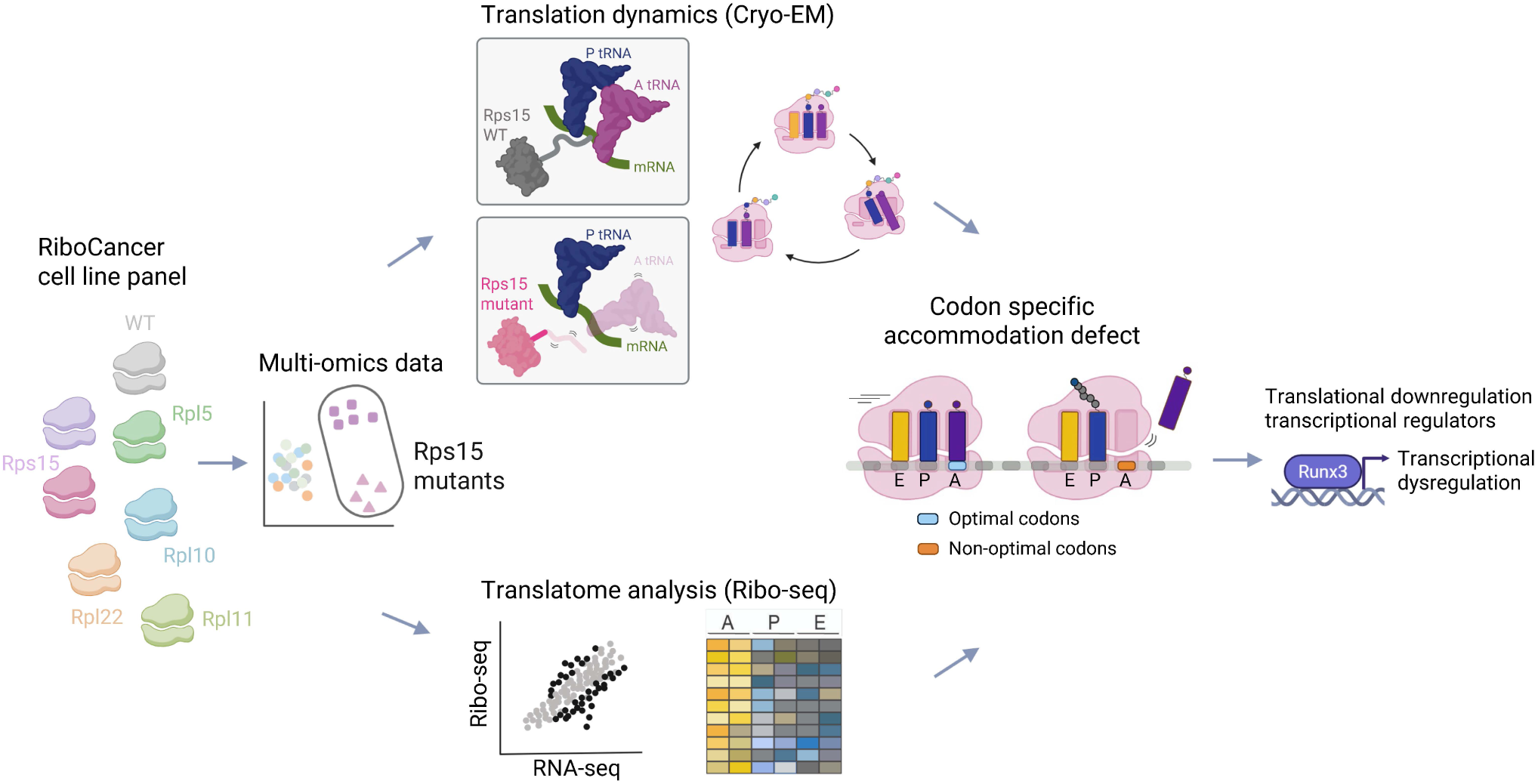

## INTRODUCTION

The ribosome is a precise molecular machine whose translation activity emerges from the coordinated interplay of its protein and RNA constituents. Accumulating evidence—from differential ribosomal protein (RP) paralog incorporation to altered RP stoichiometry and rRNA modifications—demonstrates that even subtle changes in ribosomal composition or structure can reshape its translational output.^1,2^ Despite these emerging insights, we still lack a clear understanding of how clinically observed mutations perturb ribosome function, underscoring a fundamental mechanistic question in molecular biology.

Impaired ribosome production and function resulting from mutations in RPs or ribosome assembly factors causes a series of congenital disorders known as ribosomopathies. Whereas these patients suffer from disease symptoms due to a lack of cell proliferation early in life, they have an elevated cancer risk at later age.^3^ Additionally, somatically acquired mutations in the cellular translation apparatus are relatively frequent in cancer.^4^ One of the first somatic ribosomal protein (RP) mutations that was identified is the R98S missense mutation in RPL10 (uL16), occurring in 8% of pediatric T-cell acute lymphoblastic leukemia (T-ALL) samples.^5,6^ The R98 residue in RPL10 is located at the base of an essential flexible loop at a distance of 13 Å from the peptidyl transferase center. We previously performed a genomewide translatome analysis by integrating quantitative proteomics, polysomal RNA sequencing, ribosome footprinting and total RNA sequencing data from isogenic RPL10 wild type (WT) versus R98S lymphoid cells. These analyses revealed that the RPL10 R98S mutation generates a ribosome that hypertranslates oncoproteins such as JAK and STAT proteins, BCL2 and serine/glycine synthesis enzyme PSPH, with the latter inducing metabolic addiction of RPL10 R89S cells to own serine/glycine synthesis to sustain cell proliferation. These findings provide opportunities for targeting of RPL10 R98S leukemia cells with clinically used drugs, such as BCL2 inhibitor venetoclax or antidepressant sertraline that targets serine/glycine synthesis.^7–10^

The RPL10 R98S mutation is not the only somatic RP mutation in cancer, and a whole spectrum of somatic RP mutations has been described in different solid tumor types and leukemias.^4^ Missense point mutations in RPS15 (uS19) are detected in up to 20% of relapsed chronic lymphocytic leukemia (CLL) patients. These mutations typically affect the C-terminal region of the protein which extends into the decoding center of the ribosome.^11–13^ Several of these mutant proteins have been shown to be integrated in functional ribosomes, and to impact on translation fidelity.^14^ Heterozygous somatic deletions and truncating mutations in RPL5 (uL18) and RPL11 (uL5) are present in breast cancer, glioblastoma, and melanoma, as well as in hematopoietic cancers as multiple myeloma and T-ALL.^5,15–17^ In addition, impaired RPL5 and RPL11 function is a cause of the congenital ribosomopathy Diamond Blackfan Anemia (DBA) by inactivating mutation or deletions in RPL5 or RPL11 themselves or by mutations in the RPL5 regulator HEATR3.^18–20^ RPL5 and RPL11 control the activity of key oncogenes and tumor suppressors such as c-MYC and TP53 via extra-ribosomal functions independent of protein translation regulation,^21–23^ which may explain the tumor suppressor function of these RPs. Heterozygous and even homozygous inactivating mutations and deletions of RPL22 (eL22) are frequently found in T-ALL, endometrial, colorectal and gastric cancers and have been linked to overexpression of c-MYC and of stemness factor Lin28B.^24–27^ Studies in mice and drosophila suggest that, when downregulated, RPL22 can be replaced by its paralog RPL22L1, which in turn favors translation of specific mRNAs, and thus only partially compensates for the loss of RPL22.^28,29^

Whereas we have gained novel insights in how somatic RP mutations can affect cellular translation and promote oncogenesis,^4,7,14^ previous results have been obtained in a variety of cellular backgrounds, making it difficult to compare the effects of different cancer associated ribosomal lesions. To address this limitation and gain deeper mechanistic insights, we modeled the most frequent somatic RP mutations in cancer in a collection of isogenic cell lines. To obtain these cell lines, we applied CRISPR-Cas9 engineering to genetically model the mutations exactly as they occur in patients. We subsequently characterized the impact of the introduced RP mutations on ribosomal translation function by performing an extensive analysis of the cellular translatome by ribosome profiling (Ribo-seq, or sequencing analysis of ribosome bound mRNA) and quantitative proteomics. These analyses revealed the most extensive translational rewiring by the CLL associated Rps15 P131S and Rps15 H137Y mutants. Cryo-electron microscopy (Cryo-EM) analysis revealed a higher flexibility of the C-terminal moiety of mutant Rps15, which was associated with an altered elongation cycle dynamics compared to cells harboring WT Rps15. Biochemical assays suggest that these point mutations affect the decoding and accommodation of aminoacylated tRNA at ribosomal A-site at the start of each elongation cycle. Codon site occupancy analyses on our Ribo-seq data further reinforce this concept and show how decoding and accommodation of specific codons is particularly impacted by Rps15 mutations. Altogether, our data reveal how a point mutation in ribosomal protein Rps15 can mechanically alter and thus rewire the translation process, leading to a modified translatome. Furthermore, proteins of interest displaying translational dysregulation in our isogenic models carrying Rps15 mutations were significantly enriched for transcription factors and epigenetic regulators, highlighting a previously unrecognized cross-talk between dysregulation at translational and transcriptional level in these leukemias.

## RESULTS

### A RiboCancer cell line panel as a unique resource to study cancer associated ribosome mutations

We aimed to compare the impact of several recurrent somatic and congenital RP mutations on cell behavior and on ribosomal translation function in isogenic cells that accurately model the mutations as they occur in patients. As cellular background, we opted for mouse Ba/F3 pro-B-cells, which we have previously successfully used to characterize the RPL10 R98S mutation (changes detected in Ba/F3 cells could be validated in other cell lines, leukemia patient samples and cells from a knock-in mouse).^7–9,30,31^ Ba/F3 cells are of lymphoid nature, and as such representative for several lymphoid disease entities in which somatic RP defects are detected, such as T-ALL, CLL and multiple myeloma. To avoid artifacts from overexpression of mutated RPs and/or non-accurate modeling of heterozygous RP loss with shRNA or siRNA strategies, we utilized CRISPR-Cas9 engineering to knock-in the desired RP mutations at the endogenous RP genes. Also, protein tags were avoided, as these can introduce experimental biases. We generated CRISPR-Cas9 engineered Ba/F3 clones with a heterozygous truncating mutation that causes inactivation of one allele of Rpl5, Rpl11, or Rpl22 (referred to as Rpl5^+/−^, Rpl11^+/−^ and Rpl22^+/−^). Since homozygous Rpl22 loss is viable and detected at low frequency in patients,^26,27^ we also generated Ba/F3 clones with truncating Rpl22 mutations on both alleles (Rpl22^−/−^). Furthermore, we generated Ba/F3 knock-in clones with homozygous point mutations such as the T-ALL associated Rpl10 R98S mutation,^5^ as well as Rps15 P131S and Rps15 H137Y, two of the most recurrent RPS15 missense mutations in CLL (**Figure 1A**).^11,12,14^ For each genotype, at least three independent single cell derived clones were generated (**Table S1**). Whereas mRNA and protein levels of the CRISPR-Cas9 engineered RP gene were not affected in the Rpl10 and Rps15 point mutant clones, reduction of Rpl5, Rpl11 and Rpl22 levels in the knock-out clones of these RPs was confirmed by qRT-PCR and western blotting (**Figure 1B-D**) and in our RNA-seq and proteomics datasets described below (**Supplementary Figure 1**). All RP mutations were associated with impaired growth (**Figure 1E**), which is consistent with what is observed in ribosomopathies, where RP mutations are initially associated with a cellular proliferation defect.^3^ In our Ba/F3 RiboCancer cell line panel, the growth defect was most pronounced for the Rps15 mutations and was associated with an increased fraction of cells in the G0/G1 cell cycle phases for the Rps15 P131S mutant (**Supplementary Figure 2**). To get a first impression on the impact of the introduced RP mutations on ribosomal translation performance, we evaluated nascent protein synthesis using an O-propargyl-puromycin (OPP) assay, which showed a 20-40% reduction for all RP mutants (**Figure 1F**).

**Figure 1:**
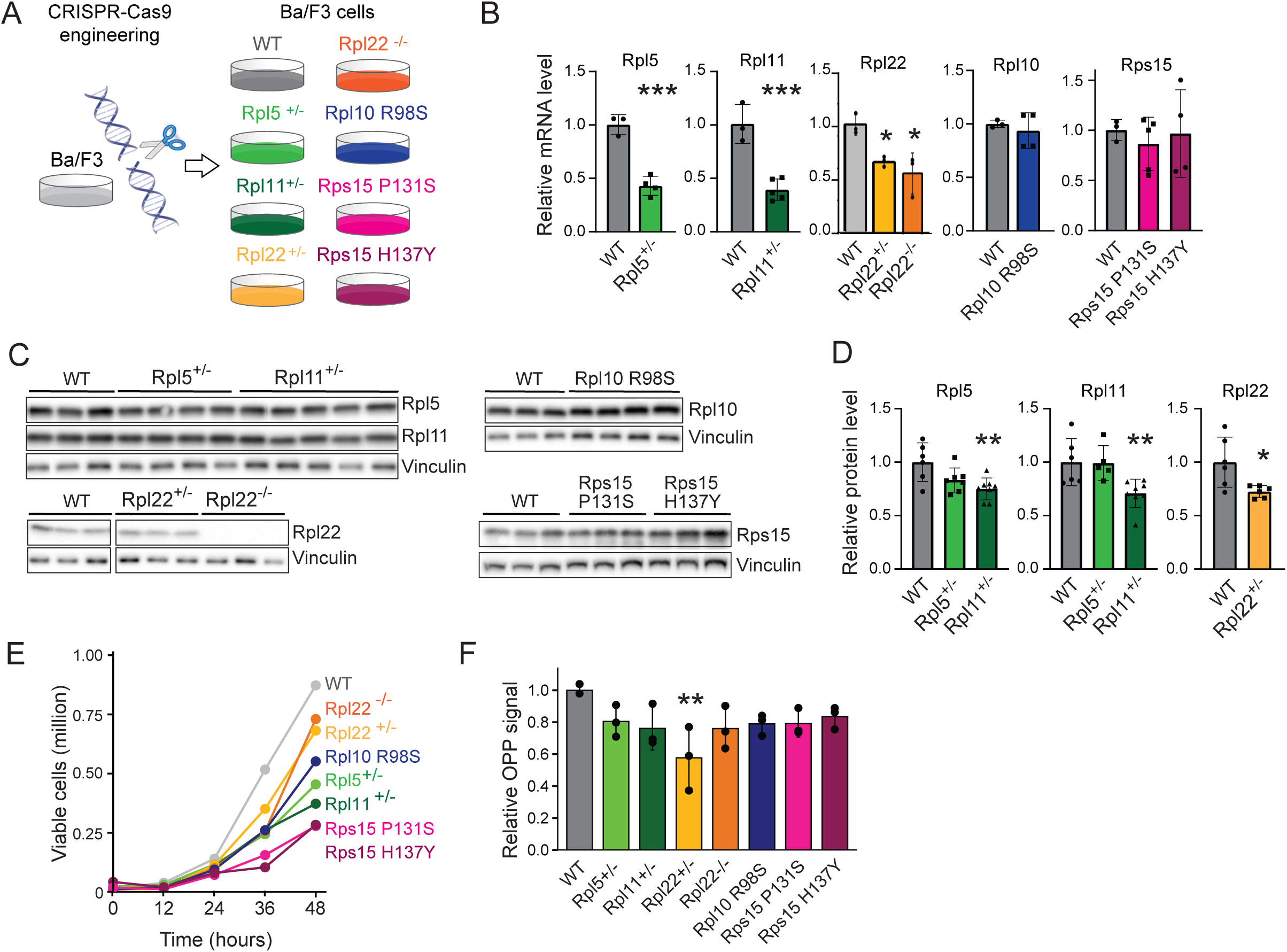
A RiboCancer cell line panel to study cancer associated ribosome mutations. **A)** Scheme illustrating the generation of the RiboCancer cell line panel. **B)** qRT-PCR analysis of the respective RP mRNA expression levels. The dots in the bar graphs represent data obtained from different single cell clones of the indicated genotype. The bars indicate mean +/− SD. **C)** Western blot analysis of the indicated RP protein expression levels. Each lane contains sample from a different single cell clone of the indicated genotype. The image shows results from a representative experiment. Vinculin served as loading control. **D)** Quantification of western blot analysis of RP protein levels. Data were pooled from 2 independent experiments, with at least 3 independent clones per genotype in each experiment (dots in the bar graphs represent data from individual clones). The bars indicate mean +/− SD. Vinculin signal was used to normalize for protein loading. **E)** Proliferation curve of single cell clones of the indicated genotypes. Each dot represents the average value of three independent clones per genotype. **F)** Nascent protein synthesis levels were evaluated by the OPP assay. Data are represented relatively to WT, dots in the bar graph represent data from independent clones of the indicated genotype. The bars indicate mean +/− SD. Statistics was done using ANOVA followed by Tukey’s HSD post hoc test. * p<0.05; ** p<0.01; *** p<0.001

### Translatome analysis reveals extensive translational dysregulation by CLL-associated Rps15 mutants

To verify whether the introduced RP mutations result in translation efficiency (TE) changes of specific transcripts, we analyzed our RiboCancer cell line panel using translatome analysis consisting of total RNA-seq, Ribo-seq quantification of ribosome protected RNA fragments (RPFs) and shotgun quantitative proteomics (**Figure 2A**). For each of these -omics experiments, at least three independent single cell clones per genotype were analyzed. In the principal component analysis (PCA) plots of these -omics datasets, the Rps15 P131S and H137Y mutants were distinct from all other analyzed genotypes (**Figure 2B**). Next, we utilized the RNA-seq and Ribo-seq data to identify genes with a significant TE change (i.e. transcripts with a differential RPF ribosome binding after normalization for changes in total transcript levels). Whereas the Rpl5^+/−^, Rpl11^+/−^, Rpl22^+/−^ and Rpl22^−/−^ cells showed little or no TE changes, it was clear that the point mutants (Rpl10 R98S, Rps15 P131S and Rps15 H137Y) showed extensive translational dysregulation (**Figure 2C**). The Rps15 P131S and H137Y mutants showed by far the highest number of TE changes, with respectively 10% and 7% of the detected transcripts showing a significant TE change. Among these, more transcripts showed a significant reduction in TE, whereas significantly increased TE was less common. Since translational dysregulation has been associated with alterations in rRNA modifications,^32^ we determined the rRNA 2’-O-Methylation profile of each RP mutant cell line by RiboMethSeq analysis. Although Rps15 mutants clustered apart from the others, only subtle differences in 2’-O-methylation were detected (**Supplementary Figure 3**). In conclusion, whereas RP knock-outs displayed limited TE changes, the RP point mutants, in particular the CLL-associated Rps15 mutants, displayed profound translatome rewiring.

**Figure 2:**
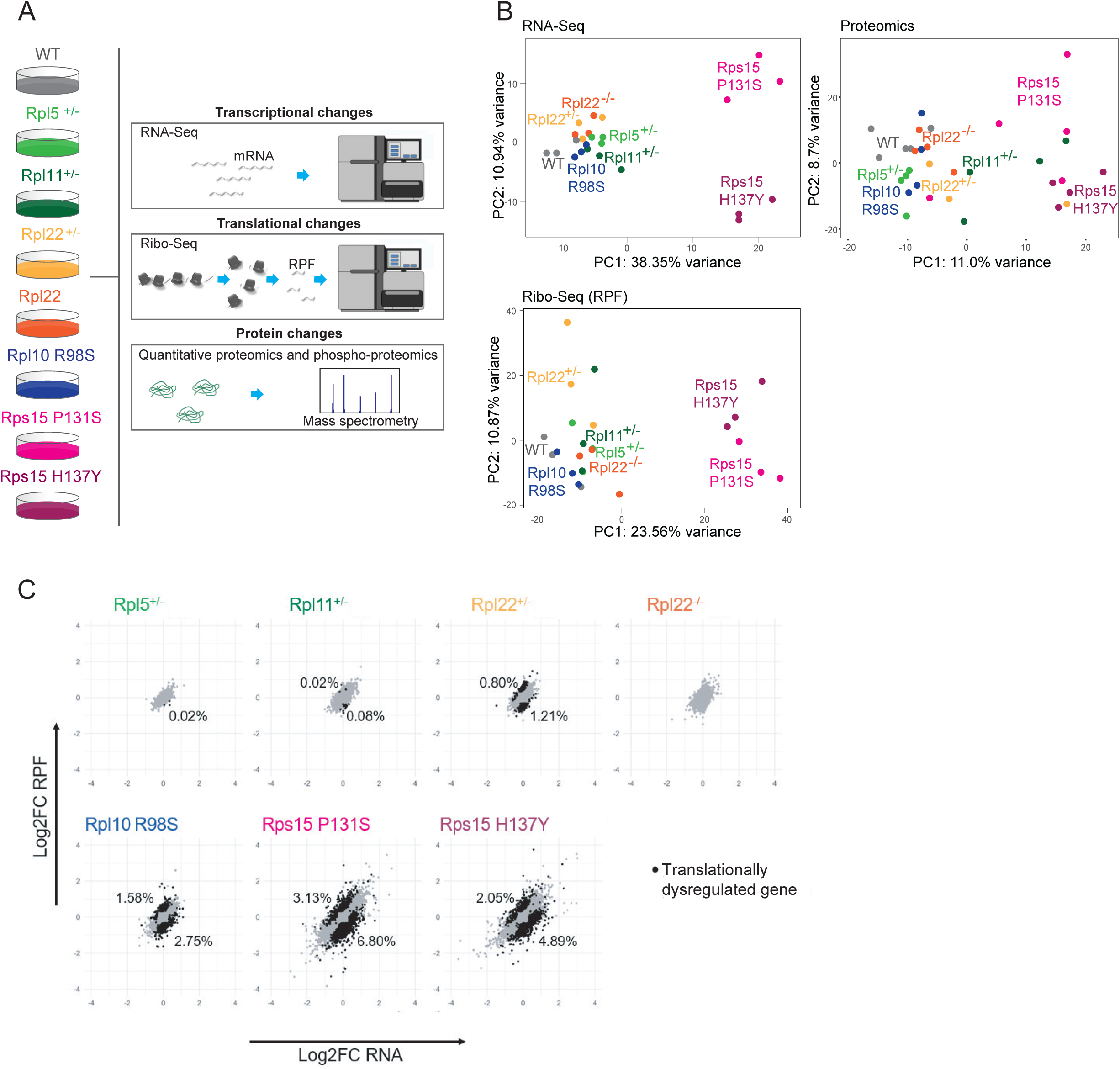
Translatome analysis reveals extensive translational dysregulation in Rps15-mutant cells. **A)** Scheme of the generated multi-omics translatome dataset. **B)** PCA plots of the total RNA-seq (RNA-seq), Ribo-seq (RPF) and quantitative proteomics dataset. Each dot in the plots represents an independent single cell derived clone of the indicated genotype. **C)** Translation efficiency (TE) analysis of RiboCancer cell line panel. Scatter plot of Log2 fold change ribosome protected fragments (Log2FC RPF - measured by Ribo-seq) versus total RNA log2 fold change (Log2FC RNA - measured by total RNA-seq) of each of the indicated RP mutants versus WT. Each dot in the plot represents a gene and shows the average value obtained from three independent clones per genotype that were analyzed. Genes with a significant TE change (= altered ribosome occupation per expressed mRNA molecule are shown in black. The percentages of detected genes that have a significantly up- or downregulated TE are indicated in the plots. These percentages correspond to the following absolute numbers: 0 Up, 3 Down (Rpl5^+/−^ vs WT); 3 Up, 12 Down (Rpl11^+/−^ vs WT); 96 Up, 145 Down (Rpl22^+/−^ vs WT); 0 Up, 0 Down (Rpl22^−/−^ vs WT); 194 Up, 337 Down (Rpl10 R98S vs WT); 371 Up, 807 Down (Rps15 P131S vs WT); 270 Up and 645 Down (Rps15 H137Y).

### Cryo-EM analysis shows elongation cycle defects for CLL-associated RPS15 mutants

In order to gain deeper insights into the mechanism driving the observed translatome rewiring for Rps15 mutants, we used cryo-EM and single particle analysis to precisely characterize the conformational changes of these ribosomes during translation.^33,34^ We determined the atomic 3D structures of translating (elongating) ribosomes purified from WT and Rps15-mutant Ba/F3 cell lines (**Figure 3A**). Translation elongation is a cyclic process governed by the transient association of elongation factors (mostly GTP-hydrolyzing proteins), which allow conformational changes of the small ribosomal subunit relative to the large one, as well as tRNA movement, peptide bond formation and translocation of the ribosome to the next codon to be decoded. This gives rise to 3D ribosomal structures that are clearly distinguishable from each other, depending on the elongation step in which the ribosome resides (**Figure 4A**). Our cryo-EM analysis allowed us to sort out ribosomal particles and determine the atomic structures of elongating ribosomes in the so-called “classical PRE-”, “rotated PRE-” and “post-translocation” (POST) steps, with resolutions ranging from 2.7 to 3.5 Å (**Supplementary Figures 4 and 5**). Comparison of the 3D structures for WT and Rps15-mutant ribosomes showed that both P131S and H137Y point mutations lead to a structural destabilization of the C-terminal part of Rps15. Indeed, while WT ribosomes displayed clearly interpretable cryo-EM densities reaching until the final C-terminal K145 residue in the classical-PRE state, and until T136 in the rotated-PRE and post-translocation (POST) states, ribosomes harboring the P131S mutation only revealed cryo-EM densities up until residues H128 for both the rotated-PRE and POST states **(Figure 3B, Supplementary Figure 6)**. Similarly, structures from the H137Y mutant ribosomes displayed density corresponding to Rps15 up to its residues G129 (rotated-PRE) and P131 (POST), respectively **(Figure 3B, Supplementary Figure 6)**. This density loss in the mutants could arise either from the partial degradation and truncation of the C-terminus of Rps15, or from an increased flexibility of the C-terminal moiety upon its mutation. Western blot analysis of polysome fractions revealed similar migration for the WT and mutant Rps15 proteins, suggesting that the mutant protein is full length within translating ribosomes **(Figure 3C),** and supporting that the point mutations lead to a structural disorganization of the Rps15 C-terminal domain. Next, to assess the dynamics of the elongation cycle, we counted the particles that populate each of the 3D structures/elongation steps: a higher number of particles represent a more stable and long-lived structural species. We reproducibly observed a decreased fraction of particles of the classical-PRE state (with three tRNAs at A, P, and E locations) and an increase of post-translocation state particles (with empty A site) for both Rps15 P131S and H137Y mutant ribosomes compared to WT (**Figure 4B**). We checked for putative bias in our processing scheme by performing AI-based structural heterogeneity assessment with cryoDRGN.^35^ This yielded similar results, with both Rps15-mutant ribosomes showing very few particles in the classical-PRE and rotated-1 PRE elongation states (**Supplementary Figure 7, Supplementary movies 1 and 2**). The classical-PRE state corresponds to the step where a new aminoacylated-tRNA has been decoded and accommodated at the ribosomal A-site. Our structural results thus point towards dysfunction of these very first steps of the elongation cycle. As we found almost no mutant ribosomes with tRNA at the A position, we reasoned that A-sites of elongating mutant ribosomes might be more available for the binding of drugs targeting this site. To validate this hypothesis, we performed ribosome collision experiments,^36^ using low doses of either anisomycin (A-site specific) or emetine (E-site specific) (**Figure 5A**).^37,38^ In the absence of drugs, the basal collision level was equally low in WT and Rps15 P131S mutant ribosomes. Whereas E-site antibiotic emetine induced similar collision levels in WT and Rps15-mutant ribosomes, A-site antibiotic anisomycin caused significantly more collision events in Rps15 P131S mutant compared to WT ribosomes (**Figure 5B-C, Supplementary Figure 8**). Of note, cellular viability assays using either emetine or anisomycin were in line with our collision experiments, with an increased sensitivity of Rps15-mutant Ba/F3 cells to anisomycin compared to WT (**Figure 5D**), which was not observed with emetine. This reinforces the idea that the A-site of Rps15-mutant ribosomes is overall more vacant (and thus more available for anisomycin accommodation) than their WT counterpart. Altogether, our cryo-EM analysis supports that Rps15 mutations alter the structural stability of the C-terminal Rps15 domain, which in turn affects the dynamics of the translation elongation cycle of these mutant ribosomes, destabilizing the decoding/ accommodation step of a new aminoacylated tRNAs at the ribosomal A-site.

**Figure 3:**
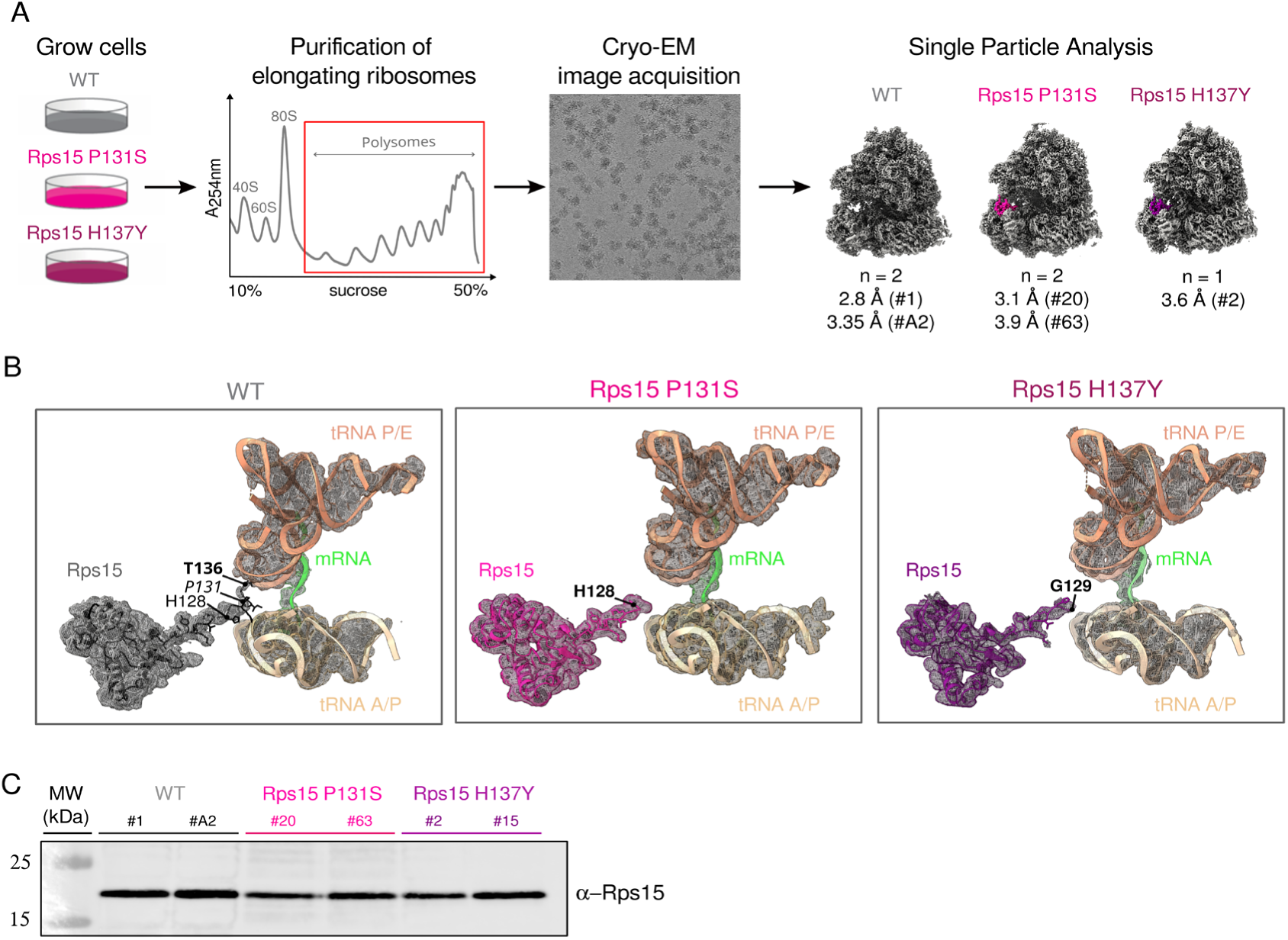
CLL-associated Rps15-mutants display increased flexibility of their C-terminus. **A)** Outline of the experimental design for the cryo-EM analysis of translating ribosomes. The indicated resolutions below the structures on the right correspond to the cryo-EM maps of Rotated-2 PRE ribosomes. After the indicated resolution, the hashtag followed by a number between brackets indicates the analyzed CRISPR-Cas9 clone. **B)** Details of the cryo-EM maps of WT (left panel), Rps15 P131S (middle panel) or Rps15 H137Y (right panel) mutant ribosomes in Rotated-2 PRE states. Cryo-EM densities are displayed in transparent mesh, while modelled atomic structures are shown with ribbons and sticks. Rps15 is shown in dark grey (WT) or magenta (Rps15 mutant), mRNA in green, tRNA in golden and brown. **C)** Western Blot analysis of Rps15 in polysomal fractions from WT, Rps15 P131S or Rps15 H137Y cells. For each genotype, the two tested clones are indicated by the number behind the hashtag.

**Figure 4:**
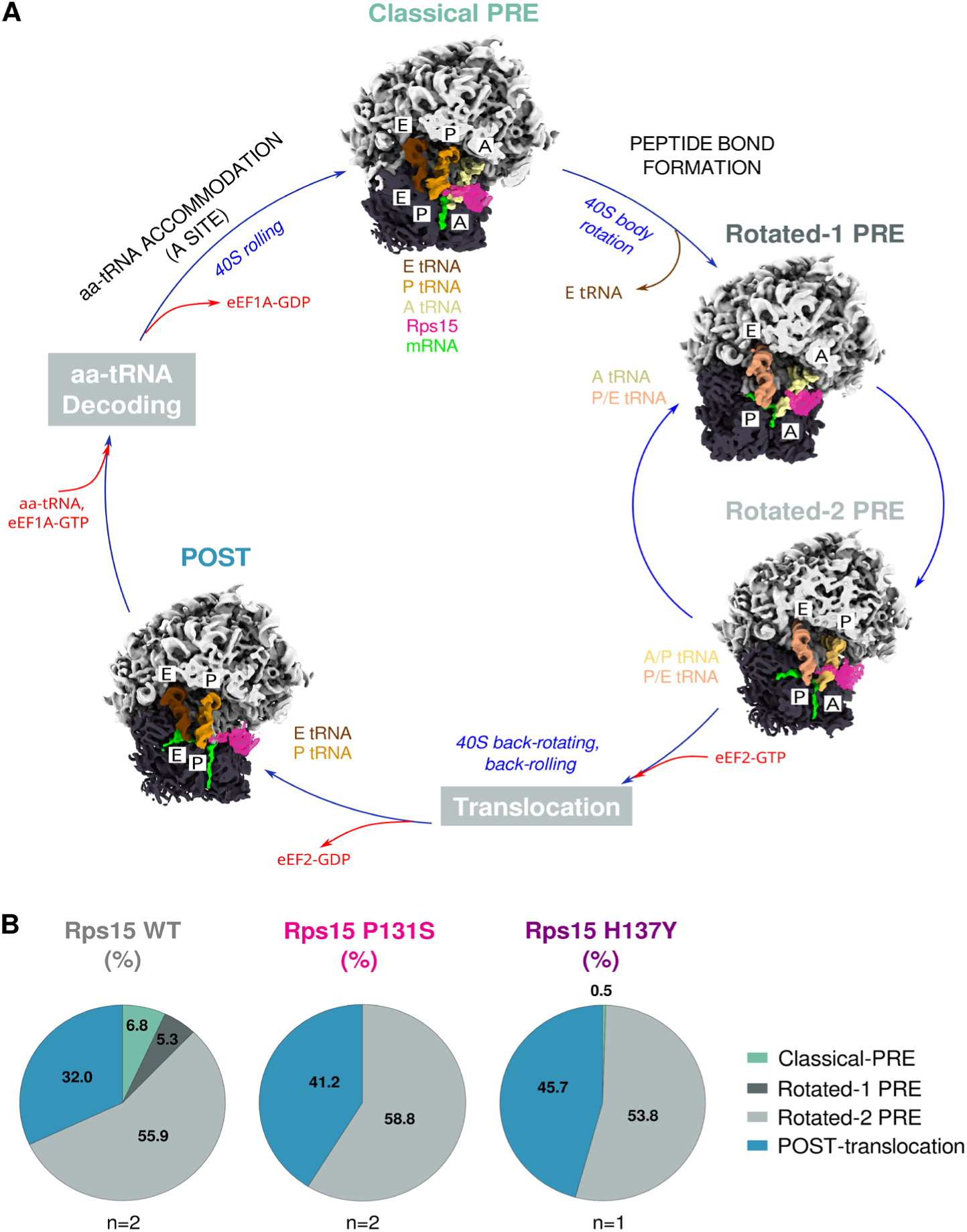
Rps15 mutations alter elongation cycle dynamics. **A)** Overview of the obtained cryo-EM maps (Rps15 WT polysomes), placed within the translation elongation cycle. Translocation and aminoacylated-tRNA decoding complexes were not experimentally observed and are represented by grey rectangles. All cryo-EM maps are filtered to 7 Å. Small and large ribosomal subunits are shown in dark and light grey, respectively; tRNAs are in shades of yellow/brown, and Rps15 is in magenta **B)** Relative abundance of ribosomal elongation complexes in all datasets. Two biological replicates (n=2) were performed for Rps15 WT and Rps15 P131S polysomes, and the percentage represents the average relative abundance. One cryo-EM experiment was performed on Rps15 H137Y polysomes (n=1). **Supplementary Figure 4** reports absolute particle numbers per experiment.

**Figure 5.**
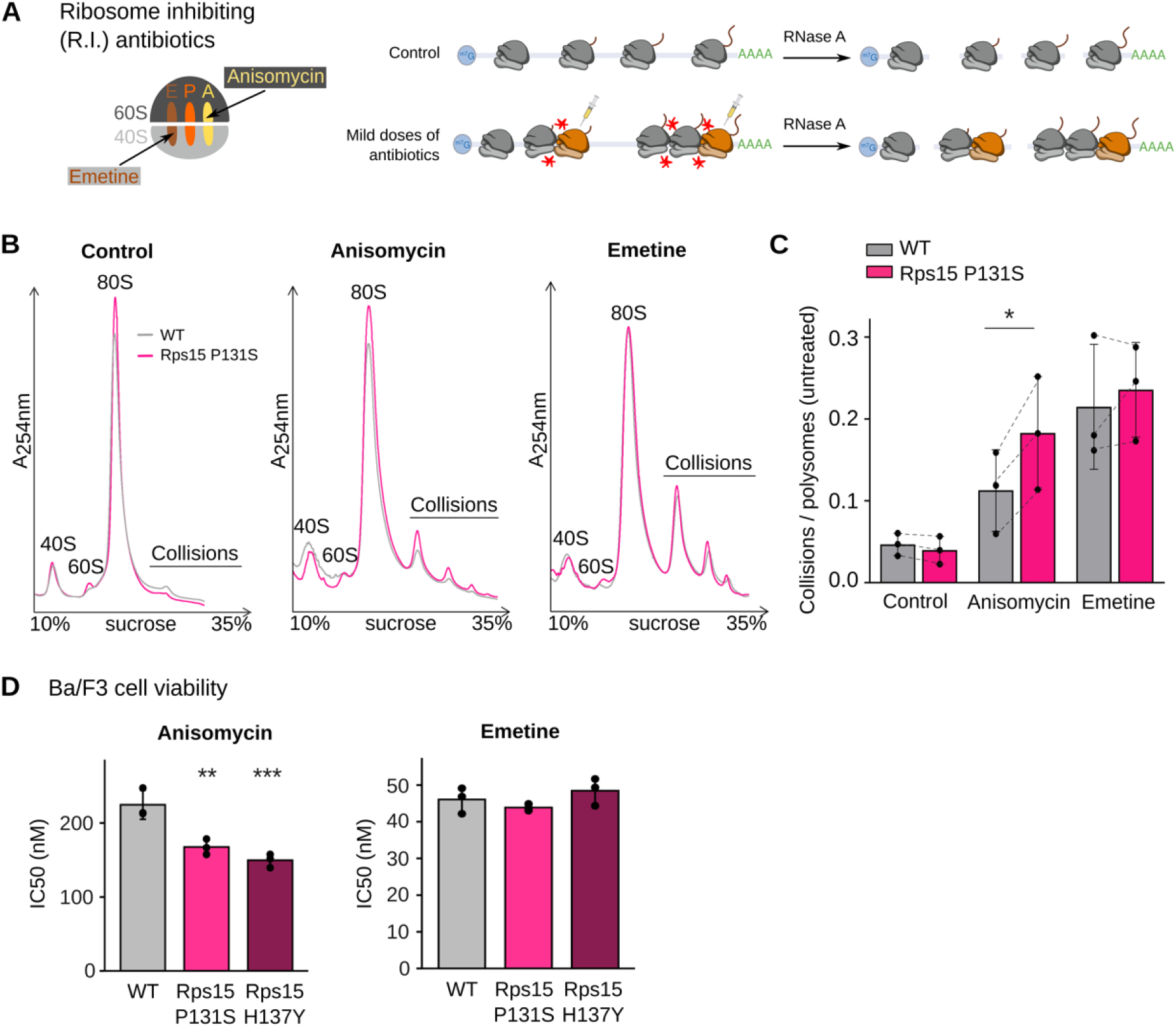
Translating Rps15-mutant ribosomes have increased vacant A-sites but not E-sites compared to WT ribosomes. **A)** Principle of ribosome collision experiments induced by ribosome inhibiting antibiotics (R.I. antibiotics) anisomycin and emetine.^36^ The syringe indicates ribosomes that have integrated the antibiotics and are therefore stalled. Ribosome collisions are indicated by red stars. The image on the left shows ribosomal tRNA binding sites targeted by the R.I. antibiotics used in this assay. **B)** Collision experiments. Polysome profiles of RNase A treated WT (grey curves) or Rps15 P131S (magenta curves) Ba/F3 cell lysates, without addition of antibiotic (control) or with incubation with anisomycin, or emetine. **C)** Quantification of the levels of collisions for each condition (control, anisomycin or emetine) performed three times on independent clones. The ratio corresponds to the area under the curve of the collision peaks, divided by the area under the curve of polysomes in the untreated condition (which are shown in **Supplementary Figure 8**). The bars represent mean +/− SD. Statistics was performed using a two-way ANOVA, followed by a Sidak’s multiple comparisons test. * p<0.05 **D)** Bar plot representing the IC₅₀ values for anisomycin and emetine treatments in WT and Rps15-mutant Ba/F3 cells. Results were obtained with an ATPlite assay and show increased sensitivity of Rps15 mutant versus WT cells to anisomycin but not emetine. The graph shows data from a representative experiment in which three clones of each genotype were assessed. Statistical significance was calculated using a two-way ANOVA, followed by a Tuckey’s HSD post hoc test. **p<0.01 and ***p<0.001

### Translational dysregulation in Rps15 mutant cells is associated with particular codons

Our data above indicates that mRNA translation dysregulation of Rps15-mutant cells is occurring at the level of accommodation of aminoacylated tRNA in the ribosomal A-site. This may be due to a generalized tRNA accommodation defect induced by the Rps15 mutations, or the defect may be specific to certain tRNAs. To investigate this, we used again our RiboSeq data. The single nucleotide resolution of these data allows to calculate the frequency (occupancy) of each codon at the ribosomal A-, P-, and E-sites, and hence to evaluate codon/tRNA-specific phenotypes.^39,40^ When comparing codon occupancy across our entire RiboCancer cell line panel, both Rps15 mutants showed a profile that was distinct from the WT and from other RP mutants in our collection, in particular in respect to A-site codon occupancy (**Figure 6A**). Next, we investigated the Rps15 mutants in more detail, and compared codon occupancy in A-, P- and E-sites of these mutants versus WT (**Figure 6B**). This showed that not all 64 codons behaved in the same way, with certain codons showing accumulation in the ribosomal A-site (codon group 1, **Figure 6B**), while other codons rather accumulated in the ribosomal P- or E-site (codon groups 3 and 4), or were unaltered (codon group 2). These results support that translational dysregulation and tRNA accommodation in Rps15 mutants show some codon-specificity.

**Figure 6:**
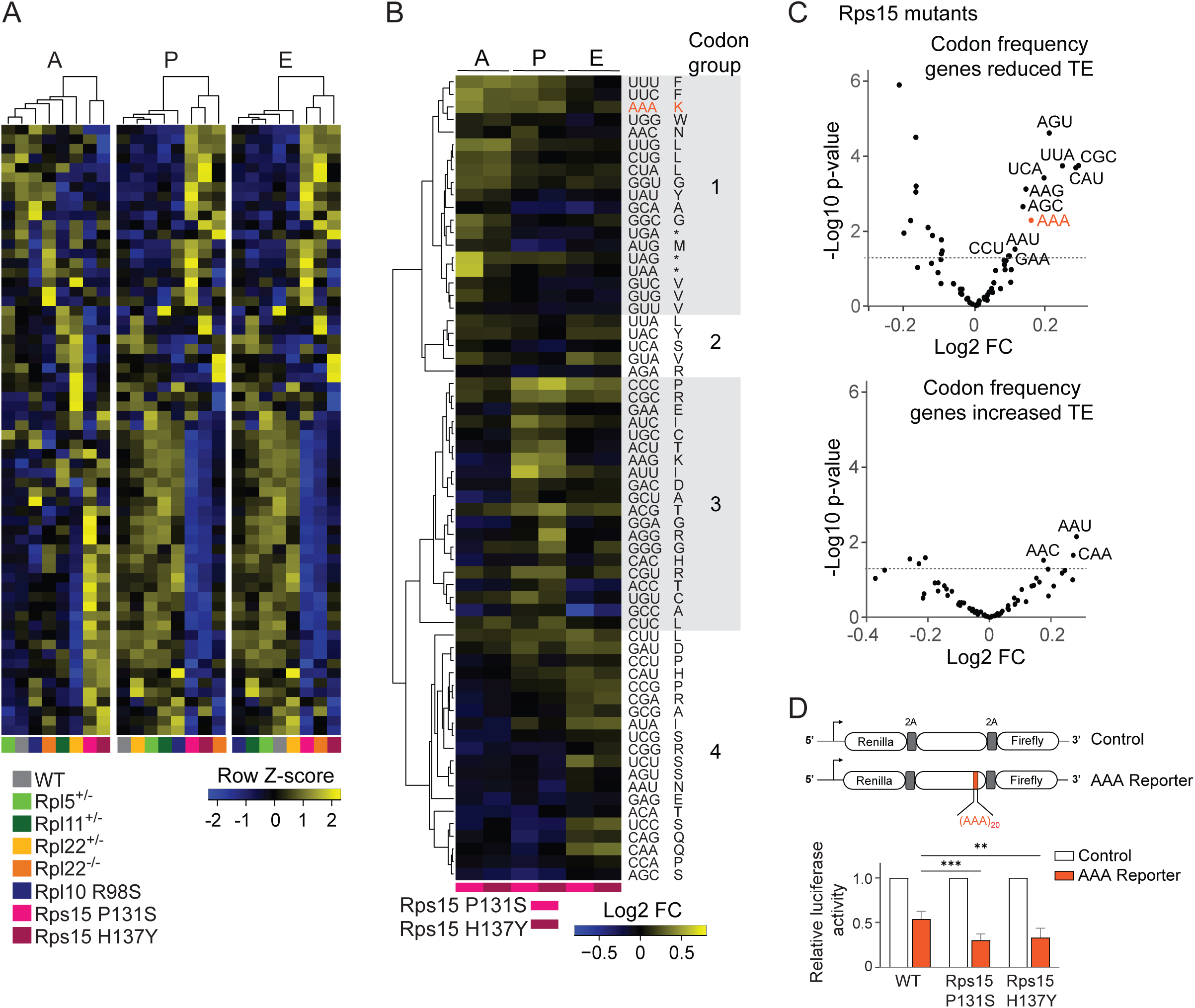
Translational dysregulation in Rps15 mutant cells shows codon specificity. **A)** Heatmap representing the row Z-score of codon occupancy for each of the 64 codons in the ribosomal A-, P-and E-sites in each genotype of our RiboCancer cell line panel. The heatmap is clustered at the codon and genotype level. This analysis is based on Ribo-seq data as explained in the methods section. **B)** Heatmap representing the Log2FC of codon occupancy in the ribosomal A-, P- and E-sites in Rps15 mutant versus WT ribosomes. The Log2FC values were determined based on Ribo-seq data as explained in the methods section. Data are clustered at the codon level, allowing identification of four distinct groups of codons (1,2,3,4) showing accumulation in the A-site (group 1), P-site (group 3), E-site (group 4) or no accumulation (group 2). **C)** Volcano plots showing the difference in codon frequency between a gene group of interest versus the mouse genome. Gene groups correspond to transcripts that are translationally downregulated (reduced TE, top panel) or upregulated (increased TE, bottom panel) in both Rps15 P131S and H137Y mutant cells relative to WT, as determined by Ribo-Seq analysis of three independent clones per genotype. Statistical significance was assessed by T-tests for each codon. The 11 codons that are significantly more frequent in genes with reduced TE in Rps15 mutants are labelled, and the AAA codon - further examined in reporter assays - is highlighted in orange. **D)** Top: Scheme illustrating the dual-luciferase reporter constructs containing either a control sequence (control) or the control sequence in which an (AAA)_20_ repeat was inserted (AAA reporter). The bar plot at the bottom represents the ratio of Firefly/Renilla signal for the Ba/F3 cell lines with the indicated genotypes. The bars represent mean +/− SD. Statistics was done on 6 replicates using a two-way ANOVA, followed by a Sidak’s multiple comparisons test. ** p<0.01; *** p<0.001

Since we identified many mRNAs with decreased TE in Rps15 mutants (**Figure 2C**), we next evaluated if certain codons are significantly overrepresented in these mRNAs, as their high frequency in these mRNAs might negatively impact TE. To this end, we compared the frequency of each codon in transcripts that were translationally downregulated (significantly reduced TE) in Rps15 mutants versus its frequency in the overall mouse transcriptome (**Figure 6C**, upper graph). This revealed 11 codons (AGU, UCA, UCU, AGC, CAU, AAG, AAA, CGC, UUA, AAU and CCU) that were significantly overrepresented in open reading frames (ORFs) of mRNAs with reduced TE. Interestingly, analysis of codon frequency in the ORFs of translationally upregulated transcripts in Rps15 mutants did not reveal such strong codon overrepresentation (**Figure 6C**, lower graph), suggesting that codon usage was mainly impacting on translational downregulation. Focusing on the 11 codons that were enriched in the translationally downregulated transcripts, our attention was attracted to the lysine codon AAA. This codon is known to slow down translation elongation when iterated.^41^ Furthermore, it was the only codon amongst the 11 identified codons that showed such a strong accumulation in the A-site of Rps15 mutants (**Figure 6B**, orange codon). Using dual-luciferase reporter assays, we experimentally validated that an mRNA containing AAA stretch was significantly less efficiently translated by Rps15 mutant ribosomes compared to WT (**Figure 6D**). Altogether, these data support that defects in the decoding / accommodation of specific codons by Rps15 mutant ribosomes negatively impact the translational efficiency of a subset of mRNAs.

### Rps15-mutant associated translational defect reduces translation efficiency of genes with a high 11-codon score linked to epigenetic and transcriptional regulation

After having identified 11 codons that are overrepresented in translationally downregulated genes in Rps15-mutant cells, we were interested in characterizing the function of genes that are enriched for these codons. First, we verified which ORFs in the mouse genome are enriched for these codons. For this purpose, we ranked all genes in the mouse genome based on occurrence of these 11 codons (weighted 11-codon score)^42^ in their ORF, and ran GSEA analysis on this ranked gene list. ORFs of genes related to nucleosome, histones, epigenetic and transcriptional regulation, cell cycle regulation and DNA repair showed the strongest overrepresentation of these 11 codons (**Figure 7A**). Interestingly, when running GSEA analysis on the genes with reduced TE in Rps15-mutant vs WT cells, the same or very similar terms linked to nucleosome assembly, epigenetic regulation, histones, transcriptional regulation and DNA repair were identified (**Figure 7B-C**). This result supports that presence of these 11 codons in the ORF is a major factor dictating TE in Rps15-mutant cells, and shows that the ORFs that are most enriched for these codons are also the ones showing the strongest translation defect in Rps15-mutant cells. When verifying the leading edge genes of the most enriched terms in our GSEA analyses, we noted that these mainly consisted of histones (**Table S2-3**). In agreement with this, many histones were reduced in TE in Rps15-mutant cells (**Figure 7D**), and ORFs encoding histones showed a 1.5 fold higher weighted 11-codon score as compared to the rest of the mouse ORFs (**Figure 7E**).

**Figure 7:**
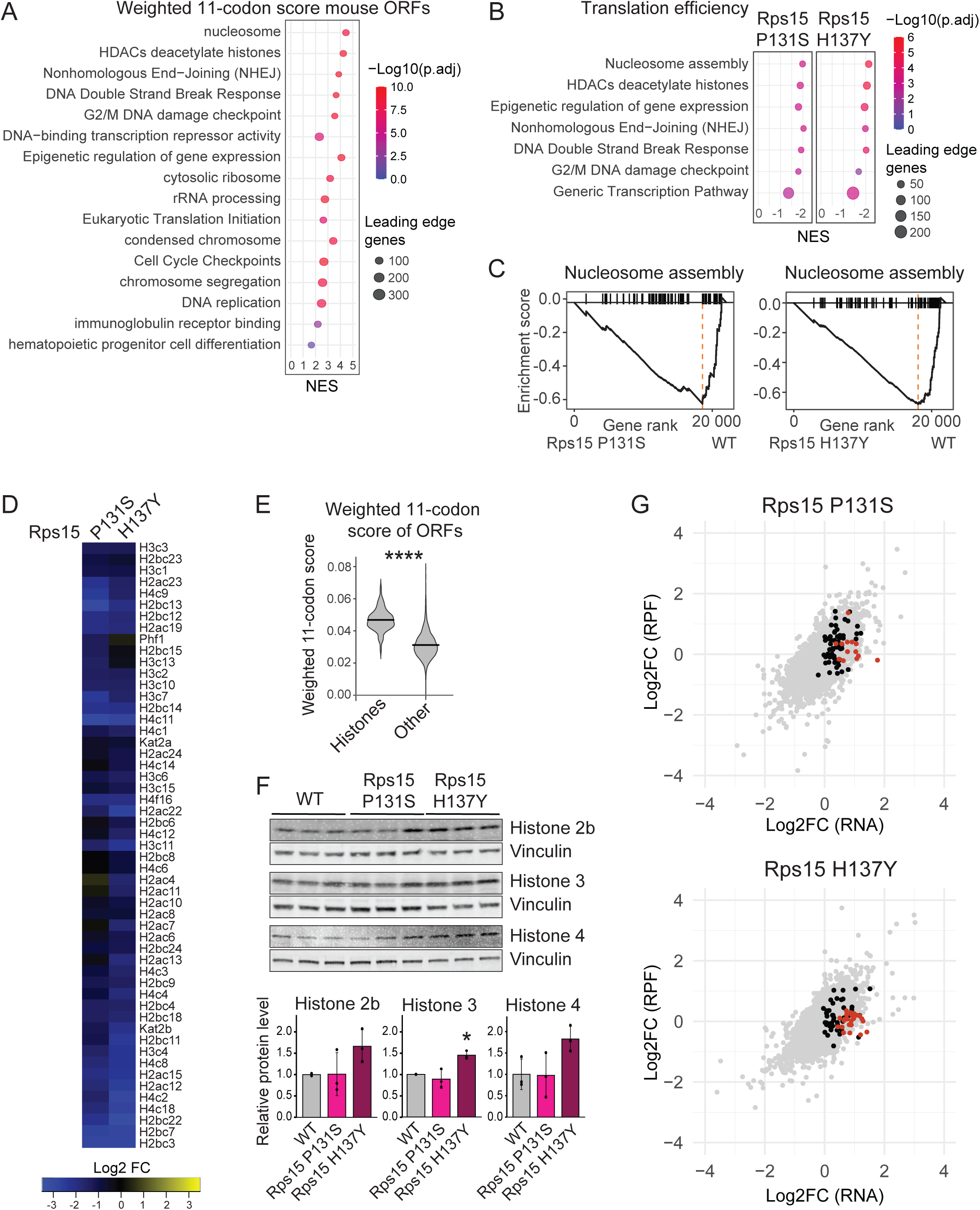
Rps15 mutant associated translational defect reduces translation efficiency of genes with a high 11-codon score linked to epigenetic and transcriptional regulation. **A)** GSEA bubble plot showing the gene sets with the strongest enrichment for the weighted 11-codon score in their ORF. NES: normalized enrichment score. **B)** GSEA bubble plot of translation efficiency (TE) illustrating the gene sets with a reduced TE in Rps15 P131S versus WT and Rps15 H137Y versus WT cells. **C)** GSEA enrichment plots showing reduced TE of genes involved in nucleosome assembly in Rps15 P131S versus WT and Rps15 H137Y versus WT cells. **D)** Heatmap illustrating the reduction in TE of many histone genes in Rps15 P131S versus WT and Rps15 H137Y versus WT cells. **E)** Violin plot comparing the weighted 11-codon score of mouse histone ORFs versus all other ORFs in the mouse genome. Statistics was done using a T-test. **** p<0.0001 **F)** Western blot analysis of histone protein levels in WT, Rps15 P131S and Rps15 H137Y Ba/F3 cells. Each lane corresponds to a different single-cell derived CRISPR-cas9 clone of the indicated genotype. The bottom part of the panel shows the quantification of the histone western blot signals, where vinculin was used as loading control to normalize the protein expression data. The bars indicate mean +/− SD. Statistics was done using a one-way ANOVA followed by a Tukey HSD post hoc test. * p<0.05 **G)** Scatter plot of translation efficiency (TE) for Rps15 P131S vs WT (upper plot) and Rps15 H137Y vs WT (lower plot). Each dot in the plot reflects the Log2FC RPF versus Log2FC RNA of a gene detected in our Ribo-seq and RNA-seq experiments, and shows the average value obtained from three independent clones per genotype that were analyzed. Histone genes that were identified as ‘buffered’ (see methods) are indicated with orange dots (n=13 for Rps15 P131S vs WT; n=25 for Rps15 H137Y vs WT). The black dots represent histone genes that were not meeting the statistical requirements to be regulated by ‘buffering’ (n=59 for Rps15 P131S vs WT; n=47 for Rps15 H137Y vs WT).

To evaluate whether the reduced TE of histone genes in Rps15-mutant cells also resulted in lower histone protein levels, we performed western blot analysis, as information on histones was missing in our quantitative proteomics data. Histone protein levels were unaltered or rather increased in Rps15-mutants as compared to WT (**Figure 7F**). Indeed, when we verified the plots of RNA-seq versus RPF counts, we noticed that a significant number of histone genes were ‘buffered genes’, and that reduced RPF counts of these genes were counteracted by increased total RNA levels^43^ (**Figure 7G**), explaining our result at the protein level.

To identify TE changes that also result in relevant protein expression changes in Rps15-mutant cells, we analyzed our shotgun proteomics results from these cells. GSEA analysis using GO molecular function confirmed that many downregulated proteins were involved in transcriptional regulation, DNA repair and cell cycle/mitotic regulation (**Figure 8A, Table S4**), similar terms as we detected in the GSEA analysis of 11-codon enrichment in the mouse transcriptome (**Figure 7A**). To identify proteomic changes that are driven by reduced translation, we integrated the proteomics data with the obtained Ribo-seq and RNA-seq data. This revealed that among the genes involved in transcriptional regulation, one of the most pronounced translationally downregulated factors confirmed at protein level was Runx3 (**Figure 8B-D**), a runt-family transcription factor regulating hemopoiesis that is dysregulated in blood cancers.^44–46^ Interestingly, the Runx3 ORF is located in the 99^th^ and 98^th^ percentile of all ORFs in the mouse genome with respect to occurrence of AGC and CGC (**Supplementary Figure 9**), two codons that are part of the 11-codon signature that is hard to translate by Rps15-mutant ribosomes. To investigate whether the Runx3 downregulation was having any functional consequences in the Rps15-mutant cells, we applied ChEA3.^47^ ChEA3 is a bioinformatic tool that allows to predict the transcription factors that are causing the detected gene expression changes in our RNA-seq data from our Rps15 P131S and H137Y versus WT cells.

**Figure 8:**
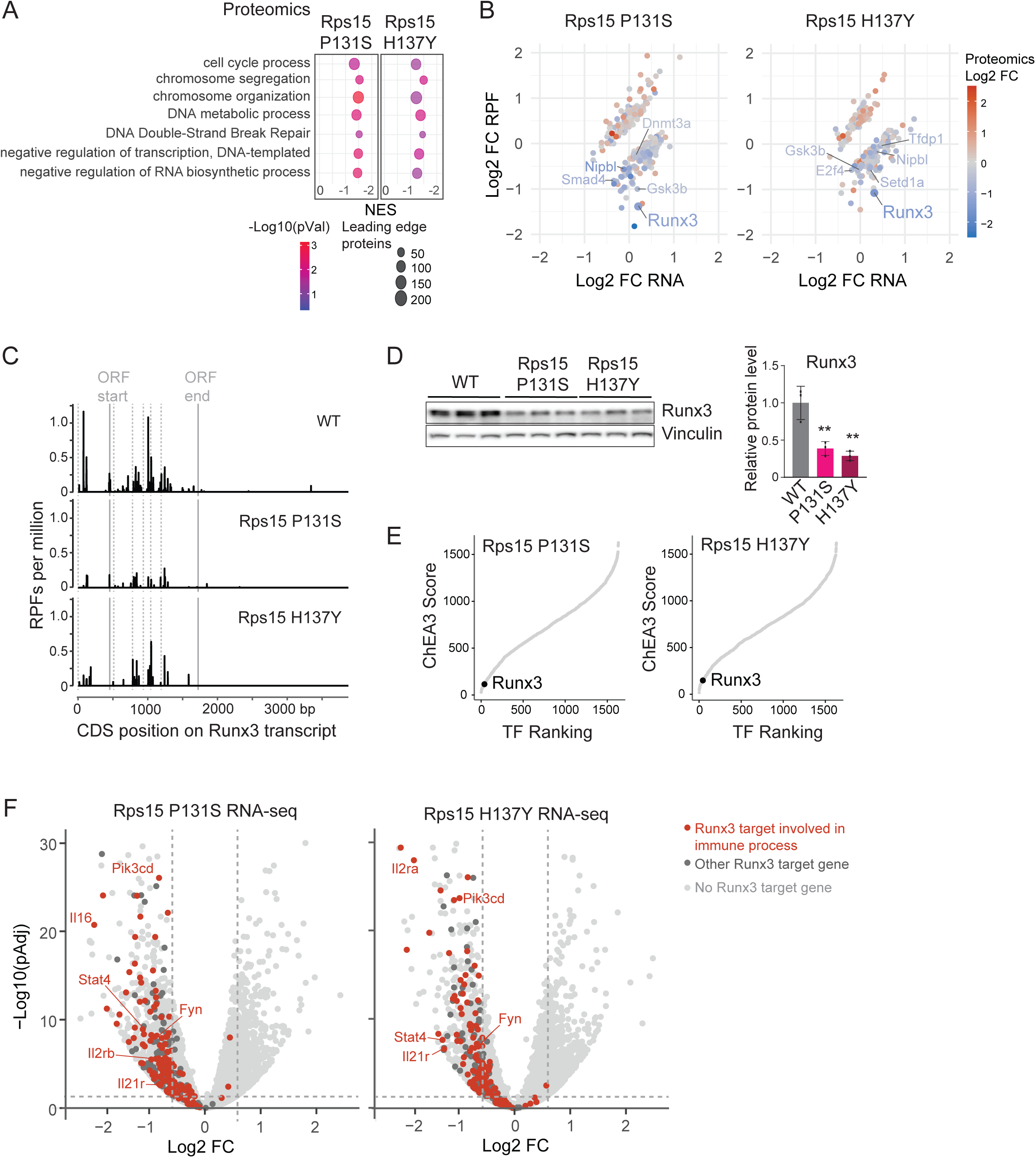
Runx3 is translationally downregulated in Rps15 mutant cells, resulting in downregulation of target genes involved in immune regulation. **A)** GSEA bubble plot of quantitative proteomics showing downregulation of proteins linked to the indicated GO molecular functions in Rps15 P131S versus WT and Rps15 H137Y versus WT cells. **B)** Scatter plot of RPF Log2FC versus total RNA-seq Log2FC of Rps15 P131S versus WT. Only genes that have a significant TE change and that could also be quantified in the quantitative proteomics analysis are displayed, and the proteomic change is color coded. Each dot in the plot represents the average value obtained from three independent clones per genotype that were analyzed. **C)** Density plot showing the distribution of RPFs covering Runx3 transcript ENSMUST00000056977 in WT and RPS15 mutant cells. The dotted grey lines indicate exon boundaries, and the continuous gray lines indicate ORF start and end sites. **D)** Western blot analysis of Runx3 expression in three independent CRISPR-Cas9 clones of the indicated genotypes. Vinculin served as loading control. The bar plot displays quantification of the Runx3 western blot signal, normalized to vinculin and to the Runx3 abundance in the WT samples. Statistics was done using a one-way ANOVA followed by a Tukey HSD post hoc test. ** p<0.01 **E)** Scatter plot of ChEA3 score versus transcription factor (TF) ranking when comparing each Rps15 mutant to WT. A lower score and rank indicate a higher predicted contribution of the transcription factor to the observed transcriptional changes detected by RNA-seq. Runx3 is highlighted and ranked 40^th^ out of 1632 evaluated transcription factors in Rps15 P131S versus WT, and 45^th^ out of 1632 for Rps15 H137Y versus WT. **F)** Volcano plot of RNA-seq results from Rps15 P131S versus WT and Rps15 H137Y versus WT cells on which Runx3 target genes identified in the ChEA3 analysis are indicated. Each dot in the plot represents a gene and shows the average value obtained from three independent clones per genotype that were analyzed. Target genes involved in immune regulation are indicated in orange, whereas Runx3 targets with another function are indicated in dark grey. A couple of example genes involved in immune regulation are labeled with their gene name.

Out of the 1632 transcription factors that were evaluated by ChEA3, Runx3 ranked respectively 40^th^ and 45^th^, indicating that Runx3 was a major transcription factor causing downregulation of its target genes in Rps15-mutant cells (**Figure 8E-F**). Interestingly, the large majority of Runx3 regulated target genes in Rps15-mutant cells had a function in immune signaling and immune regulation (**Figure 8F, Supplementary Figure 10**). Altogether, these findings indicate that among the genes with reduced TE in Rps15 mutants, many are linked to a specific codon frequency signature, which is characteristic of genes involved in nucleic acid binding and transcriptional regulation. The Runx3 transcription factor is strongly translationally downregulated in Rps15-mutant cells, causing misexpression of target genes involved in immune regulation.

## DISCUSSION

Ribosomes have long been viewed as “blind protein factories,” assumed to possess identical composition and production capacity in every cell of an organism. However, numerous findings have nuanced this monolithic view. It is now well established that heterogeneity in ribosomal proteins and RNAs can influence the cellular translatome, giving rise to the concept of “specialized ribosomes”.^1,48^ In recent years, somatic mutations have been found to recurrently target RPs, implicating them in cancer development and progression.^49^ Whereas independent studies have described translational phenotypes for some of these RP mutants,^7,9,14^ differences in the design of experimental models and utilized cellular backgrounds preclude comparison of the impact of the different RP mutations on ribosome function. In this regard, the RiboCancer cell line panel that we constructed, modeling the most recurrent cancer associated RP mutants in an isogenic background, is a unique resource to mechanistically compare these mutations. The RP mutations were knocked-in at the endogenous gene locus, without introducing any protein tags, ensuring accurate modeling of the mutations as they occur in cancer samples. Furthermore, Cas9 was only transiently expressed during the genome engineering, avoiding off-target effects. In all of our -omics studies, at least three independent single cell derived clones per genotype were analyzed, and results from the different clones of each genotype were remarkably consistent as can be appreciated in the PCA plots of our -omics analyses, supporting that off-target effects in our clone collection are minimal. In line with many other studies in literature^50–52^, all modeled RP mutations in our RiboCancer panel were associated with a global mRNA translation and proliferation defect. However, when looking at TE changes, it was clear that two distinct classes of RP mutations emerged in our collection: the RP knock-out models displayed minimal TE changes, versus the RP point mutants in which up to 10% of genes showed a significant TE change (most often a reduced TE). This observation suggests a difference in the oncogenic role of these two classes of RP mutations, and suggests that dysregulation of extra-ribosomal functions may be of major importance for the inactivating RP defects that we studied. Indeed, RPL5 and RPL11 have well characterized extra-ribosomal functions in regulating notorious oncogenes and tumor suppressors such as TP53 and c-MYC.^21–23^ Also for RPL22, extra ribosomal regulation of TP53 via MDM4 splicing was recently described.^53^

In our translatome screen, the most pronounced TE changes were detected for the CLL-associated Rps15 mutants. We detected more genes with a reduced TE in these mutants, which aligns with Ribo-seq that has been performed on CLL patient samples.^54^ Using cryo-EM, we further characterized the mechanistic basis of the translation defects in these Rps15 mutants. Rps15 is a protein of the small ribosomal subunit whose eukaryotic-specific C-terminal domain interacts with A- and P-site tRNAs, as well as mRNA.^55,56^ Crosslinking experiments and structural studies previously suggested that this C-terminal domain plays a key role in the accommodation of aminoacyl-tRNAs at the A-site during elongation.^57–59^ A recent cryo-EM analysis revealed that specific residues of Rps15 engage in hydrophobic and hydrogen-bond interactions with A- and P-site tRNAs, while its terminal residues contact the +4 position of the mRNA (ie, the first nucleotide of the A-site mRNA codon) to stabilize codon–anticodon pairing. This C-terminal region functions as a “two hinges and a latch” system, coordinating P-site tRNA movements and A-site tRNA accommodation.^13^ Our cryo-EM and biochemical analyses of elongating ribosomes harboring either wild type or mutant Rps15 support these findings. Despite proper incorporation of Rps15 into ribosomes, point mutations (P131S, H137Y) increase the flexibility of the Rps15 C-terminus, likely weakening interactions with incoming aminoacyl-tRNAs and impairing their decoding or accommodation at the A-site. This defect may lead to ribosome accumulation in the post-translocation state, as observed for the Rps15 mutants. Although overall translation remains functional, we nonetheless observed a reduced activity as well as codon-specific defects, particularly at AAA (Lys) codons for Rps15-mutant ribosomes. These results align with previous studies implicating Rps15 in facilitating the accommodation of charged Lys, Glu, and Arg codons.^56^ We propose that CLL-associated C-terminal mutations disrupt this regulatory role, introducing mechanical “glitches” in translation elongation that reshape the translatome of the mutant cells. Our Ribo-Seq and proteomics analysis suggest that translation of mRNAs encoding charged nucleotide binding proteins will be primarily affected by CLL-related Rps15 mutations.

Interestingly, we identified 11 codons that are strongly enriched in ORFs of genes that are translationally downregulated in Rps15-mutant cells. When screening the genome for ORFs with the highest occurrence of these 11 codons, the highest scoring ORFs were encoding genes involved in regulation of chromatin structure, transcription and DNA repair, such as histone genes. These were precisely the gene classes that we found to be translationally downregulated in Rps15-mutant cells, supporting that frequency of these codons in an ORF is dictating TE in these mutants. Nevertheless, other factors besides codon usage also seem to be relevant for these mutants. In this regard, we also found a negative correlation of TE with GC-content and ORF length for Rps15 H137Y but not Rps15 P131S (**Supplementary Figure 11**). This underscores that although Rps15 H137Y and P131S show similar phenotypes in our cellular assays and in the -omics and cryo-EM analyses, there also are functional differences. These differences have also been suggested by the results of Bretones et al.,^14^ and are of interest for further exploration.

Our results show that Rps15 mutations are associated with profound translational dysregulation of factors involved in regulating chromatin structure and gene transcription. Whereas cancer research has focused a lot on studying transcriptional dysregulation in the past, these results reveal defects in ribosomes as a previously unrecognized cause of downstream transcriptional dysregulation. This can have profound cellular effects, as we could show downregulation of transcription factor Runx3, resulting in dysregulated expression of many Runx3 target genes involved in immune regulation. RUNX3 is an important transcription factor in hematopoietic development that has been described to be misexpressed in hematopoietic disorders such as acute myeloid leukemia (AML) and CLL.^44–46^ Our findings invite to investigate the role of RUNX3 misexpression in CLL in more detail.

Rps15-mutant Ba/F3 cells are hypo-proliferative, with an increased doubling time and reduced translation activity compared to wild type Ba/F3 cells, modeling the hypoproliferative phase of ribosomopathies.^3^ In particular in the context of the T-ALL associated RPL10 R98S mutation, we previously showed that secondary mutations are required to compensate this slow growth phenotype and transition to the hyperproliferative phase.^30^ Also in CLL patients, presence of RP mutations as RPS15 is associated with a higher mutation burden, suggesting that also RPS15 defects require secondary compensatory mutations.^30^ As we observed profound transcriptional rewiring downstream of the Rps15 mutations, we speculate that Rps15 defects induce a premalignant transcriptional state, and that additional transcription factor defects may be needed to turn this into a malignant transcriptional state. The co-occurrence of mutations in RP or ribosome biogenesis-related genes with mutations in the TP53 transcription factor have been found in numerous blood and solid cancers,^60,61^ and TP53 mutations are among the strongest prognostic markers in CLL.^62^ Therefore, we suspect that the effects of RPS15 mutations, seemingly innocuous at first, may cause selective pressure to acquire compensatory mutations in genes that are key for CLL and cancer development such as TP53, ATM or NOTCH.

Overall, this study demonstrates that frequent somatic RP mutations impose distinct translational rewiring, offers unprecedented mechanistic insight into CLL-associated RPS15 mutations, and reveals an unexpected cross-talk between translational and transcriptional dysregulation in RP-mutant cells.

## METHODS

### Cell Lines

Ba/F3 cells were obtained from German Collection of Microorganisms and Cell Cultures GmbH (DSMZ; ACC 300) and were grown in RPMI 1640 medium (Gibco) supplemented with 10% fetal calf serum (FCS, Gibco) and with 1 ng/mL interleukin-3 (IL-3, Miltenyi Biotec). Cultured cells were regularly confirmed to be mycoplasma negative.

### CRISPR-Cas9 engineering

A design for sgRNAs targeting the RP genes of interest was made using the Synthego CRISPR Design Tool (https://design.synthego.com/). The designed sgRNA sequences (**Table S5**) were cloned into pSpCas9(BB)-2A-GFP (pX458, Addgene #48138). One million Ba/F3 cells were electroporated in 2 mm cuvettes with 10 µg of pX458 containing sgRNA in 100 µL serum-free RPMI 1640 using 6 square wave pulses (175V, 0.1 ms interval) followed by transfer to 2 mL of pre-warmed recovery medium (RPMI 1640 supplemented with 10% FBS, 1 ng/mL IL-3, 1mM Sodium Pyruvate (ThermoFisher) and 1 mM MEM Non-Essential Amino Acids (ThermoFisher)). At 24 hrs after electroporation, GFP-positive cells were sorted on a BD FACS Aria III, and plated at a density of one cell per well in tissue-culture treated 96 well plates (Greiner Bio-One) in RPMI 1640 supplemented with 10% FBS, 1 ng/mL IL-3 and 100 µg/mL primocin (InvivoGen).

In case of knock-in models of point mutations, the following adaptations were done to the procedure described above: a 117-nt donor oligo (Integrated DNA Technologies) containing the mutant allele of interest was added in the Ba/F3 electroporation at a concentration of 1 µM (**Table S5**), and 10 µM SCR7 (Sigma-Aldrich) was added to the recovery medium. At 24 hours after electroporation, GFP-positive cells were sorted and plated in ClonaCell-TCS Medium (Stemcell Technologies) supplemented with 10% FBS, 1 ng/mL IL-3 and 100 µg/mL primocin at a density of 1000 - 8000 cells/well in 6-well plates. Once visible by eye, single cell colonies were picked and transferred to liquid cultures in RPMI 1640 supplemented with 10% FBS, 1 ng/mL IL-3 and 100 µg/mL primocin. Growing single cell derived clones were further expanded in RPMI 1640 supplemented with 10% FBS, 1 ng/mL IL-3 and analyzed by Sanger sequencing on genomic DNA to verify the desired genetic RP modification and to exclude clones containing modification(s) of *in silico* predicted off-target effects. Cell clones that have undergone the whole CRISPR-Cas9 engineering procedure as described above but that were unmodified at the genomic loci of interest, were used as wild type (WT) control cell clones.

### qRT-PCR

RNA was extracted with the RNeasy mini kit (Qiagen) and reverse transcribed with the GoScript Reverse Transcription System (Promega). qRT-PCR reactions were performed using GoTaq qPCR Master Mix (Promega) on a CFX Connect Real-Time PCR Detection System (Bio-Rad), using primers as listed in **Table S6**. Data were analyzed using the ΔΔCT method, using Hprt or Actb as reference gene.

### Immunoblotting

Cells were lysed in lysis buffer (Cell Signaling Technology) with addition of cOmplete protease inhibitors and 5mM Na3VO4 (Roche). Proteins were separated on 4-15% Criterion TGX Precast protein gels (Bio-Rad) and transferred to PVDF membranes using a Pierce G2 Fast Blotter (Thermo Scientific). Primary antibodies used for immunodetections are listed in **Table S7** Secondary antibodies allowing chemiluminescent or fluorescent detection were used and blots were imaged on an Azure C600 (Azure Biosystems). Blot intensities were quantified using ImageJ software.

### Cell cycle analysis

One million of WT or Rps15-mutant cells were fixed in absolute ethanol and stored at −20 °C overnight. Cells were then rinsed with PBS and centrifuged at 600 g for 10 min. The pellet was resuspended for 15 min at RT with a PBS staining solution containing 1 µg/mL DAPI [Sigma, D9542] and 0.05 % Triton X100. Cells were centrifuged at 1,200 g for 10 min and the pellet was resuspended in 1 mL PBS to remove excess of DAPI. Samples were analyzed for DNA content using a CytoFLEX S Flow Cytometer and the CytExpert software.

### O-propargyl-puromycin (OPP) assay

For OPP labeling, we used the Click-IT OPP Alexa Fluor 647 Protein synthesis Assay kit (Thermofisher). A number of 100 000 cells were seeded in 96-well plates in culture medium supplemented with 10 uM OPP. After 15 min, cells were fixed in 4% formaldehyde for 15 min, and permeabilized for 15 min in PBS 0.5% Triton X-100. Cells were then incubated for 30 min with the Click-IT OPP reaction cocktail prior to rinsing with the Click-IT reaction rinse buffer. Cells were finally resuspended in PBS and analyzed on a MacsQuant X (Miltenyi Biotec) flow cytometer.

### RiboMethSeq

We profiled rRNA 2’-O-methylation by RiboMethSeq^63^ at 76 positions. The 76 high confidence positions corresponding to conserved sites with human rRNA for which a guide C/D box snoRNA has been ascribed were selected based on sequence comparison between mouse, human and yeast rRNA sequences, mouse 2’-O-Me data from the snOPY database (http://snoopy.med.miyazaki-u.ac.jp/snorna_db.cgi?mode=sno_search&organism=Mus_musculus) and 2Ome-seq data from Incarnato *et al.*.^64^ Briefly, the presence of the methyl group at the 2’ position protects the phosphodiester bond located at the 3’ of the methylated site from alkaline hydrolysis. Thus, the presence of 2’-O-me at a given nucleotide *n* induces a decrease in RNA hydrolysis at the *n +* 1 position. The 2’-O-Me level is calculated by comparing the fragment end-count at each position with the 6 nt on either side (C-score). RiboMethSeq was performed as previously described^65^ using Illumina technology on the RibosOMICS and PGC / PBGT platforms of the CRCL starting from 100 ng of total RNA of each cell line. After sequencing, adapter sequences were trimmed off using Trimmomatic,^66^ extracted reads were aligned on the mouse rRNA sequences (5.8S rRNA: NR_003280.2 ; 28S rRNA: NR_003279.1 ; 18S rRNA: NR_003278.3) using Bowtie 2, and 5ʹ and/or 3ʹ end reads were counted at each rRNA nucleotide position (https://github.com/RibosomeCRCL/ribomethseq-nf). Using the rRMSAnalyzer package (https://github.com/RibosomeCRCL/rRMSAnalyzer), the quality control was performed to identify putative outliers, C-scores were computed (flanking window of ±6 nucleotides), and data visualization was drawn.

### Ribo-Seq and RNA-seq of total RNA

For each genotype, three independent single-cell derived clones were analyzed. For each clone, 10-50 million cells were treated for 1 min with 0.1 mg/mL cycloheximide, followed by washing of cells in 10 mL ice-cold PBS supplemented with 0.1 mg/mL cycloheximide, pelleting of cells and flash freezing of the cell pellet in liquid nitrogen. Cell pellets were shipped to OHMX.bio (Evergem, Belgium) for further processing. Briefly, cell pellets were lysed in 750 μl of ice-cold Lysis buffer supplemented with 100 mg/mL cycloheximide. After debris removal, the lysate was divided for total RNA purification and for subsequent footprinting.

For RNA-seq of total RNA, RNA was purified from the lysate with the RNA Clean and Concentrator-5 kit (Zymo Research), followed by NEBnext polyA mRNA magnetic isolation and processing with the NEBNext Ultra II directional RNA library prep kit (NEB) in combination with NEBNext Multiplex Oligos for Illumina (Dual Index Primer pairs). Libraries were sequenced using on a NovaSeq6000 platform, in PE150 mode.

For footprinting, 10 μg of lysate was digested for 45 min with RNase I at 21°C, and the reaction was stopped by adding SuperaseIn RNase Inhibitor (Thermofisher). Digested lysate was purified on a MicroSpin S-400 HR Sephacryl column (GE Healthcare), and flow through was purified with the RNA Clean and Concentrator-5 (ZymoResearch). Ribosome protected fragments (RPFs) corresponding to 28-30 nt were excised after gel electrophoresis on a Novex TBE-Urea gel (15%) (Thermofisher), and excised gel fragments were incubated overnight in molecular grade water supplemented with ammonium acetate at 4°C. The slurry was transferred to CoStar filter tubes and centrifuged, and flow through was incubated at −80°C for 2 hrs with glycogen and isopropanol. Following centrifugation, the pellet was washed with ice-cold 80% EtOH, dried and resuspended in molecular grade water. RPFs were subsequently 3’ dephosphorylated with T4 PNK (New England Biolabs) and purified using the Oligo Clean and Concentrator kit (ZymoResearch). The 3’ adapter was ligated and fragments were purified using the Small RNA Library prep kit (Lexogen). After 5’ phosphorylation of the RPFs and purification with the Oligo Clean and Concentrator kit, 5’ adapter ligation and first strand cDNA synthesis were performed according to the Small RNA Library prep kit manual. Ribosomal RNA (rRNA) was depleted by adding LNA probes from the QIAseq FastSelect rRNA HMR kit (Qiagen) during the cDNA synthesis reaction. In the indexing PCR amplification step (Small RNA Library prep kit), index primers (Lexogen) were added at both ends of the fragments. Library products were purified with AMPure XP beads, and adapter dimers and excess index primers were removed by running the samples on a Novex TBE gel (8%) (Thermofisher), the gel part corresponding to the correct library products was excised and purified. Quality of the final libraries was determined on the Agilent Bioanalyzer with the High Sensitivity DNA kit. Sequencing was performed on the Illumina Nextseq 500 system, high output v2 kit in SE75 configuration. SeqPurge (v2.0.1) was used for trimming adapter sequence from total RNA-seq reads. For Ribo-seq, adaptor sequence ‘TGGAATTCTCGGGTGCC’ was clipped. Sequences were trimmed based on sequencing quality in all datasets with the FASTX toolset (0.0.14). Sequences shorter than 20 nucleotides were hereby discarded and the quality threshold was set at 28. All contaminant PhiX, rRNA, snRNA, sn-o-RNA and tRNA sequences were filtered out. Pre-processed trimmed reads from total RNA-seq and Ribo-Seq were aligned using the nf-core/rnaseq (v3.0)^67^ to the Mus musculus (GRCm38, release 102) reference genome using STAR aligner (v.2.6.1d) and quantification was performed by Salmon (v.1.4.0) using the star_salmon option.^68^

Differential expression analysis was performed on the triplicates of two groups (WT and mutant) using the package DESeq2 (v.1.30)^69^ in the R environment.

### Purification of translating ribosomes for cryo-EM analysis

WT or Rps15-mutant Ba/F3 cells (200 ×10^6^ cells each) grown to 1.5 - 1.8 × 10^6^ cells/mL as described above were pelleted by mild centrifugation (5 min, 300 g, 4°C), followed by resuspension and lysis in lysis buffer (20 mM HEPES-KOH [pH 7.6] ; 2.5 mM MgCl_2_ ; 100 mM NaCl ; 0,25 % sodium deoxycholate [Sigma-Aldrich, D6750] ; 0,5 % IGEPAL [Sigma-Aldrich, I3021] ; EDTA-free Protease inhibitor cocktail [Roche Diagnostics, 05056489001]). Cell debris was removed by two successive centrifugations (10 min; 1,600 g; 4 °C, followed by 10 min; 10,000 g; 4 °C), after which the supernatant was kept. The resulting cytoplasmic extracts were deposited onto 10% - 50% W/V sucrose gradients in Buffer A (20 mM HEPES-KOH ; 150 mM NaCl ; 10 mM MgCl_2_ ; 1 mM DTT) and ultracentrifuged using a Beckmann Coulter SW41 rotor at 40,000 rpm for 4 °C for 2 h. Polysome profiles were read at A_254_ and collected using a Fraction Collector system (Foxy R1, Teledyne Isco). Polysome fractions were pooled, mixed with Buffer A to a volume of 11 mL, and sucrose was removed by ultracentrifugation using a Beckmann Coulter SW41 rotor at 39,000 rpm for 4 °C for 1h 10 min. Polysome pellets were resuspended in 40-50 µL of Buffer A and RNA concentration was assessed by Nanodrop measurements.

### Cryo-EM sample preparation and single particle analysis

Cryo-EM grids were prepared and imaged at METI, the Toulouse cryo-EM facility. Purified polysomes were diluted in Buffer A to reach an RNA concentration of 300-400 ng/µL. 3.5 µL of solution was deposited onto freshly glow-discharged (Pelco Easy Glow system operating at 35 mA, 30sec) holey carbon grids (QUANTIFOIL R2/1, 300 mesh with a 2-nm continuous layer of carbon on top). Grids were plunge-frozen using a Leica EM-GP automat; temperature and humidity level of the loading chamber were maintained at 18 °C and 95%, respectively. Excess of solution was blotted with a Whatman filter paper no.1 for 1.7–2.1 s and grids were immediately plunged into liquid ethane (−184 °C). Cryo-EM images were recorded on a Talos Arctica electron microscope (FEI, ThermoFisher Scientific) operating at 200 kV and equipped with a Gatan K2 summit-BioQuantum direct electron detector (counting mode). Automatic image acquisition was performed with EPU, at a magnification corresponding to a calibrated pixel size of 1.02 Å and a total electron dose of 40 *e*^−^/Å^2^ over 40 frames (maximum 1.05 e^−^/Å^2^/frame). Nominal defoci values ranged from −0.6 to −2.7 µm.

For each imaged sample – corresponding to polysomes purified from two different clones of Ba/F3 WT, two clones of Ba/F3 Rps15 P131S and one clone of Ba/F3 Rps15 H137Y – more than 7,000 stacks of frames were collected (**Supplementary Figures 4-5** and **Table S8)**. Image analysis was performed using Relion 4.0.^70^ An overview of the single particle processing scheme is presented in **Supplementary Figures 4-5**. Frame stacks were aligned to correct for beam-induced motion using MOTIONCOR2.^71^ Contrast Transfer Function (CTF) and defocus estimation were performed on the realigned stacks using CTFFIND4.^72^ The best micrographs were selected for further analysis upon CTF maximum resolution on their power spectra and visual inspection. Particles were automatically picked, then extracted in boxes of 384 × 384 pixels, using the LogPicker autopick option from RELION 4.0. A 2D classification was performed on particle images binned by a factor of 4 to sort out bad particles and free 40S subunits from complete 80S ribosomes. A subset of the latter was used for an *ab initio* 3D reconstruction, and the resulting 3D map was used as a reference for 3D classification in six classes of the particle dataset. Particles belonging to 3D classes with the best level of detail were grouped, re-extracted without imposing any binning factor, and a consensus 3D map was obtained using RELION’s 3D auto-refine option, with resolution reaching 4.0 Å and beyond depending on the sample imaged. Local masks around ribosomal A, P and E sites were designed using Chimera.^73^ Masked classification was then performed using the masked consensus 3D map as reference, which allowed to sort out ribosomes in either rotated-1 or rotated-2 PRE-translocation states, or classes of ribosomes containing at least two tRNAs, in P and E conformations and some blurred density in the tRNA A region, as well as ribosomes with poorly defined 40S subunits. A further 2D classification of the latter revealed that they corresponded mostly to free 60S subunits (**Supplementary Figure 4**). Particles belonging to 3D classes with P/P and E/E tRNAs were pooled and further submitted to a 3D classification against a low-pass filtered version of a human ribosome in a classical-PRE conformation (PDB-6Y0G,^13^), which allowed to separate classical-PRE from post-translocation ribosomes. Particles from each class were submitted to RELION’s 3D auto-refine option, followed by CTF Refinement, particle polishing and a final round of orientation refinement (3D auto-refine option, with an ad-hoc binarized and soft-edged mask). Map resolutions were estimated at this step according to gold-standard Fourier Shell Correlation (FSC) at 0.143. Post-processed maps of each ribosomal state were calculated either using the post-processing option in RELION, or DeepEMhancer.^74^

### AI-based structural heterogeneity assessment within the cryo-EM datasets

In order to characterize their nature and level of structural heterogeneity, particles belonging to the 3D consensus map of ribosomes purified from each clone type were also subjected to AI-based sorting by cryoDRGN version 2.3.0 ^75^ (**Supplementary Figure 7**). All models were trained for 50 epochs, parallelized through several GPUs. Particles were downsampled to a box size of 128 pixels with a pixel size of 3.06 Å/pixel for evaluation. We trained a cryoDRGN model on these downsampled particles with three layers and 512 nodes per layer for both the encoder and decoder network architectures (termed 512 × 3) and an 8-dimensional latent variable Z. This resulted in separated clusters of Z values, suggesting that the heterogeneity in the dataset arises from discrete particle populations with distinct compositions, consistent with the 3D classification results obtained in RELION. Particles from each cluster (**Supplementary Figure 7A**) were filtered and counted using k-means clustering implemented into cryoDRGN Jupyter notebook (**Supplementary Figure 7B**). Using cryoDRGN, we generated reconstructions corresponding to directions along the top principal component of the latent space, visualized in **Supplementary Movies 1** (Rps15 WT) and **2** (Rps15 H137Y).

### Cryo-EM maps interpretation

Atomic models and cryo-EM maps of mammalian ribosomes in classical-PRE (PDB-6Y0G, ^13^), Rotated-1 PRE (emd-2904, ^33^), Rotated-2 PRE (PDB-6Y57, ^13^) or post-translocation (PDB-5AJ0, ^33^) ribosomes were first fitted within the experimentally obtained cryo-EM maps as rigid body using the “fit” command in UCSF Chimera.^73^ Manual refinements and adjustments, as well as flexible and jiggle fittings were then realized on various chains of the models in Coot.^76^ Final atomic models were refined using REFMAC5 and Phenix.^77,78^ Final model evaluation was done with MolProbity.^79^ Overfitting statistics were calculated by a random displacement of atoms in the model, followed by a refinement against one of the half-maps in REFMAC5, and Fourier shell correlation curves were calculated between the volume from the atomic model and each of the half-maps in REFMAC5 (**Table S8**). Maps and models visualization was done with Coot, UCSF Chimera and ChimeraX.^73^ Figures and movies were created using ChimeraX and the open source video editor kdenlive (https://kdenlive.org).

### Ribosome collision assays

Cells grown to 1.5 - 1.8 × 10^6^ cells/mL were supplemented with anisomycin (Sigma-Aldrich, A9789) or emetine (Sigma-Aldrich, E2375) for 15 min at concentrations of 3.8 µM and 1.4 µM respectively. Following treatment, cells were collected by gentle centrifugation (5 min, 300 g, 4°C), and the pellet (corresponding to a total number of cells of 45 × 10^6^) was rinsed once with 10 mL of PBS and then lysed by resuspension for 20 min in 800 µL of ice-cold lysis buffer (20 mM HEPES-KOH [pH 7.6] ; 2.5 mM MgCl_2_ ; 100 mM NaCl ; 0,25 % sodium deoxycholate [Sigma-Aldrich, D6750] ; 0,5 % IGEPAL [Sigma-Aldrich, I3021] ; EDTA-free Protease inhibitor cocktail [Roche Diagnostics, 05056489001]). Cytoplasmic extracts were prepared by two successive centrifugations of 10 min at 1,600 g and 4 °C, then 10 min at 10,000 g and 4 °C, where the supernatant was kept. Extracts containing 250 µg of total RNA (as estimated from Nanodrop NP80 IMPLEN measurement) were treated with RNase A (Thermo Fisher Scientific, EN0531) at 4 mg/L for 15 min at 4°C, which was then stopped by 200 U of SUPERaseIn (Thermo Fisher Scientific, AM2694). Untreated and RNAse-digested extracts were run through 10% - 50% or 10% – 35% sucrose gradients (20 mM HEPES-KOH ; 150 mM NaCl ; 10 mM MgCl_2_ ; 1 mM DTT) respectively using a Beckmann Coulter SW41 rotor at 40,000 rpm for 4 °C for 2 h. The sedimentation profiles were read at A_254_ and collected using a Fraction Collector system (Foxy R1, Teledyne Isco). Quantifications were performed by measuring the area under the curve of polysomal fractions for untreated polysomes and collided ribosomes, using the Fityk software.^80^

### Cell viability assay

Ba/F3 cells were seeded in 50 µL RPMI 1640 medium (10 % FBS, 1 ng/mL IL3) at 5,000 cells/well in 384-well plates and treated with increased concentrations of anisomycin and emetine. Antibiotics were added to the cells using a D300e Digital Dispenser (Tecan). Cell proliferation was assessed after 48 hours using ATPlite 1step Assay system (PerkinElmer) and luminescence was measured on a VICTOR multilabel plate reader X4 (PerkinElmer).

Relative proliferation was calculated by subtracting the baseline luminescence values measured prior to treatment from those recorded after 48 hours. Data were further normalized to account for variations in initial cell number and expressed as a percentage relative to untreated cells.

### Dual luciferase reporter assays

Luciferase reporter constructs were generated by engineering modified versions of the pmGFP-P2A-K0-P2A-RFP and pmGFP-P2A-K(AAA)20-P2A-RFP plasmids.^81^ Briefly, PCR-amplified fragments encoding Renilla and Firefly luciferases were inserted into the pCI-Neo vector (Promega) using the *NheI–XhoI* and *XbaI–NotI* restriction sites, respectively. Fragments containing the stalling reporters, either lacking or containing 20 consecutive AAA lysine codons and insulated by P2A sequences, were PCR-amplified from *pmGFP-P2A-K0-P2A-RFP* and *pmGFP-P2A-K(AAA)20-P2A-RFP* (Addgene plasmids #105686 and #105688) and subsequently cloned into the above vector at the *XhoI–XbaI* sites. One million of WT or Rps15-mutant cells were electroporated using a BioRad gene pulser Xcell Eukaryotic system device [BioRad, 1652661] (square wave ; 175 V ; 6 x 2.5 ms pulse ; 0.1 ms interval) with 1 µg of control or codon reporter constructs. Electroporated cells were plated in 2 mL of recovery media (Opti-MEM + GlutaMAX [Gibco] ; 5 % FBS ; 1.25 ng/mL IL-3) in 12-well plates. Cells were lysed 24 hrs after transfection, and luciferase activities were measured with the Dual-Luciferase Reporter Assay System (Promega) in a GloMax 20/20 luminometer (Promega). The RLuc activity was normalized against the activity of Fluc.

### Translation efficiency calculation

After mapping and quantifying both RPFs and mRNA reads, the TE for each gene was estimated as the ratio of RPF-count to RNA-count. Using the wild-type (WT) condition as control, a log₂ fold-change was calculated for each gene separately from the RNA data and the RPF data. These changes (RNA and RPF) were then combined via the deltaTE methodology^43^ to detect significant changes in TE. Gene considered as “differentially translated” with increased or decreased translation were identified using different thresholds including a significant (pAdj < 0.1) difference of 25% (up or down) in TE FC as well as at least a significant (pAdj < 0.05) difference of 25% in RNA or RPF FC. Gene considered as “buffered” were identified with method described in^43^.

### Codon occupancy analysis

Reads within the 28–30 nucleotide range were extracted using the SeqKit tool (v2.1.0) and aligned to the *Mus musculus* coding transcriptome (GRCm37 – M28) with the STAR aligner (v2.7.9a). Only reads showing a perfect match were retained for downstream analyses. The optimal P-site offset was determined based on the 5ʹ ends of the selected reads relative to the translation initiation site. Specifically, the offset was inferred from the global distribution of reads around the start codon (AUG, encoding methionine), with the most frequent position defining the P-site offset. Following the methodology described in^39^, reads in the 0 and −1 reading frames were used to establish an offset of 6 nucleotides from the 5ʹ end.

Once the positions of the three ribosomal sites (E, P, and A) were identified, the reads were trimmed at the 5ʹ end to retain only these sites along with their 3ʹ flanking regions. The flanking regions correspond to the three codons downstream of the A site that are not involved in tRNA binding and serve as non-translated controls for codon occupancy estimation. Ribosome occupancy was quantified by calculating the frequency of each codon within the E, P, and A sites and in the flanking regions. The final ribosome occupancy metric was obtained by dividing the codon frequency at a given ribosomal site by the average codon frequency observed in the flanking regions.

### Analysis of codon frequency enrichment in ORFs

Codon frequency analysis was performed using open reading frame (ORF) sequences corresponding to the most abundantly expressed transcript variant of each gene in Rps15 mutant cells, as determined by Salmon.^82^ ORF sequences for these transcripts were retrieved from Ensembl. Codon frequencies for all 64 codons were obtained by dividing the count of each codon by the total number of codons in the corresponding ORF. The weighted 11-codon score for each transcript was calculated by summing, across all 11 selected codons, the product of each codon’s frequency and its FC enrichment in the group of down-translated transcripts in Rps15 mutants.^42^

### GSEA analysis

Gene sets were ordered by corresponding ranking metrics (Log2FC TE, Log2FC protein abundance or weighted 11-codon score) prior to GSEA analysis^83^ using R package ClusterProfiler^84^ with default settings. Gene Ontology^85^, Kyoto Encyclopedia of Genes and Genomes (KEGG)^86^, Reactome^87^ and MsigDB^83^ were used as annotation databases. Results were filtered on Benjamini-Hochberg calculated adjusted p-values (pAdj<0.05) or on non-corrected p-values (p<0.05) for proteomic analysis. Complete lists of enriched results are shown in **Table S2**, **S3 and S4**. To reduce redundancy in the list of significantly enriched terms that are in common between both analyzed Rps15 mutant versus WT, a Jaccard similarity matrix was calculated for each term based on core leading edge genes, followed by hierarchical clustering from which 5 to 10 representative clusters were extracted. Representative enriched terms were manually selected from the wider clusters for graphical representation.

### Transcription factor enrichment analysis

The ChEA3 R package^47^ was used for transcription factor enrichment analysis, starting from significantly downregulated genes (pAdj < 0.05 and FC < 0.67).

### Proteomics analysis of Ba/F3 cells

At least three independent Ba/F3 cell clones of each analyzed genotype (**Table S1**) were starved of serum and interleukin 3 (IL3) in RPMI 1640 medium (Gibco) for two hrs at 37°C (5% CO2), at a density of 1×10^6^ cells/mL, followed by a 30 min stimulation with 1 ng/mL IL3. After this, cells were pelleted, snap-frozen and stored at – 80 °C for later processing by the VIB Proteomics Core (Ghent, Belgium). Cell pellets were lysed in an urea lysis buffer (9 M urea, 20 mM HEPES pH 8.0 and PhosSTOP phosphatase inhibitor cocktail (Roche, 1 tablet/10 mL buffer)). Samples were sonicated with 3 pulses of 15 s at an amplitude of 20 % using a 3 mm probe, with incubation on ice for 1 min between pulses. After centrifugation for 15 min at 20,000 g at room temperature (RT), 5 mM dithiothreitol was added followed by incubation for 30 min at 55 °C. Iodoacetamide (10 mM) was added followed by an incubation for 15 min at RT in the dark. Based on a Bradford assay (Bio-Rad), 2 mg protein from each sample was used to continue the protocol. Samples were diluted with 20 mM HEPES pH 8.0 to a final urea concentration of 4 M and proteins were digested with 20 µg LysC (Wako) (1/100, w/w) for 4 hours at 37 °C. Samples were again diluted to 2 M urea and digested with 20 µg trypsin (Promega) (1/100, w/w) overnight at 37 °C. After addition of 1% trifluoroacetic acid (TFA) and 15 min incubation on ice, samples were centrifuged for 15 min at 1,780 g at RT. Next, peptides were purified on SampliQ SPE C18 cartridges (500 mg, Agilent). Columns were first washed with 5 mL 100% acetonitrile (ACN) and pre-equilibrated with 15 mL of solvent A (0.1% TFA in water/ACN (98:2, v/v)) before samples were loaded on the column. After peptide binding, the column was washed with 5 mL of solvent A and peptides were eluted twice with 700 µL elution buffer (0.1% TFA in water/ACN (20:80, v/v)). A volume of 100 µL of eluted peptides was dried completely in a SpeedVac vacuum concentrator and redissolved in 20 µL solvent A, and peptide concentration was determined on a Lunatic spectrophotometer (Unchained Labs). 2 µg of each sample was injected for LC-MS/MS analysis on an Ultimate 3000 RSLCnano system in-line connected to an Orbitrap Fusion Lumos mass spectrometer (Thermo) equipped with a pneu-Nimbus dual ion source (Phoenix S&T). Trapping was performed at 10 μL/min for 4 min in solvent A on a 20 mm trapping column (made in-house, 100 μm internal diameter (I.D.), 5 μm beads, C18 Reprosil-HD, Dr. Maisch, Germany) and the sample was loaded on a 200 cm micro pillar array column (PharmaFluidics) with C18-endcapped functionality mounted in the Ultimate 3000’s column oven at 50 °C. A fused silica PicoTip emitter (10 µm inner diameter) (New Objective) was connected to the µPAC outlet union. Peptides were eluted by a non-linear increase from 5 to 55% MS solvent B (0.1% FA in water/ACN (2:8, v/v)) over 145 min, first at a flow rate of 750 nL/min, then at 300 nL/min, followed by a 10-min wash reaching 99 % MS solvent B and re-equilibration with MS solvent A (0.1% FA in water). The mass spectrometer was operated in data-dependent mode, automatically switching between MS and MS/MS acquisition. Full-scan MS spectra (300-1500 m/z) were acquired in 3 s cycles at a resolution of 120,000 in the Orbitrap analyzer after accumulation to a target AGC value of 200,000 with a maximum injection time of 250 ms. The precursor ions were filtered for charge states (2-7 required), dynamic exclusion (60 s; ±10 ppm window) and intensity (minimal intensity of 5E3). The fragments were analyzed in the Ion Trap Analyzer at normal scan rate. Data from LC-MS/MS runs were together using the MaxQuant algorithm (version 1.6.11.0) with mainly default search settings, including a false discovery rate set at 1% on PSM, peptide and protein level. Spectra were searched against the mouse protein sequences in the Uniprot Reference Proteome database (database release version of June 2019), containing 22,282 sequences (www.uniprot.org).

### Data analysis and statistical testing

Data analyses were mainly performed using the R statistical programming language (R Core Team, 2021. R: A Language and Environment for Statistical Computing. R Foundation for Statistical Computing, Vienna, Austria; https://www.R-project.org/). Unless otherwise specified, adjusted p-values were calculated using two-way ANOVA followed by Tukey’s HSD post hoc test (padj < 0.05: *, padj < 0.01: **, padj < 0.001: ***).

Ribosome collisions and dual luciferase reporter assays were statistically analyzed using GraphPad Prism (v10.0.3). Quantitative data are presented as mean +/− standard deviation (SD). Exact sample sizes, statistical test methods, and p-values are specified in the figure legends. A significance level of p < 0.05 was considered statistically significant.

## Data availability

RNA-seq and Ribo-seq data are available in Gene Expression Omnibus (GEO) under accession GSE310057 including Fastq files, raw counts and tables reporting differential analysis of total RNA, RPFs and TE changes when comparing mutant and WT samples.

Data deposition of RiboMethSeq data is in progress. Shotgun proteomics data from the Ba/F3 RiboCancer cell line panel have been deposited at the ProteomeXchange Consortium via the PRIDE partner repository (dataset identifier PXD051831).

Cryo-EM maps and molecular models of mouse ribosomes generated in this study have been deposited to the Electron Microscopy Data Bank (EMDB) and the Protein Data Bank (PDB), respectively, with the following accession codes: WT, Classical-PRE translocation ribosome: EMD-53473, PDB-9QZP ; WT, Rotated-1 PRE: EMD-53333, PDB-9QSA ; WT, Rotated-2 PRE: EMD-53310, PDB-9QQP ; WT, Post-translocation: EMD-53262, PDB- 9QOH ; Rps15 P131S, Rotated-2 PRE: EMD-53427, PDB-9QWT ; Rps15 P131S, Post-translocation: EMD-53307, PDB-9QQL.

## ACKNOWLEDGEMENTS

This research was funded by research grants from the European Research Council (ERC) (RIBOCANCER, 334946), FWO (G092620), Stichting Tegen Kanker (F/2024/2554) and iBOF (iBOF/23/014) to KDK, by an EHA junior research grant to MC, and by fundraising initiatives from BenEVIet and VZW Baief. SV was recipient of an FWO PhD fellowship (1S49817N), and KDK was recipient of a Collen-Francqui research professor mandate. This work was supported by the CNRS, the University of Toulouse, the Institut National du Cancer (‘ONCORIB’ Project PLBIO-2020-091), the Agence Nationale de la Recherche (ANR ANR-22-CE11-0017-01 / CRYOMETH), and the French Ligue Nationale Contre le Cancer. AA was recipient of a PhD fellowship from the Ligue Nationale Contre le Cancer. ProteoToul (JM and CF) is funded by the French Ministry of Research with the Investissement d’Avenir Infrastructures Nationales en Biologie et Santé program (ProFI, Proteomics French Infrastructure project, ANR-10-INBS-08 & ANR-24-INBS-0015).

We would like to thank the staff of the platforms that have not been included as coauthors: PGC Platform (CRCL, Lyon, France: Dir Q Wang), PBGT Platform (CRCL, Lyon, France: Dir A Ferrari), RibosOMICS Platform (CRCL, Lyon, France: Dir. F Bourdelais), METi Platform (CBI, Toulouse, Dir S Balor ; member of the national infrastructure France-BioImaging supported by the French National Research Agency ANR-10-INBS-04), CALMIP HPC facility (Toulouse, France: Dir JL Estivalezes) and the VIB Proteomics Core (Ghent, Belgium).

