## Supplementary data for "Translatome and translation dynamics analysis of a RiboCancer cell line panel reveals that leukemia-associated Rps15 mutations rewire translation through codon-specific tRNA accommodation defects"

### SUPPLEMENTARY MATERIAL

#### Supplementary figures

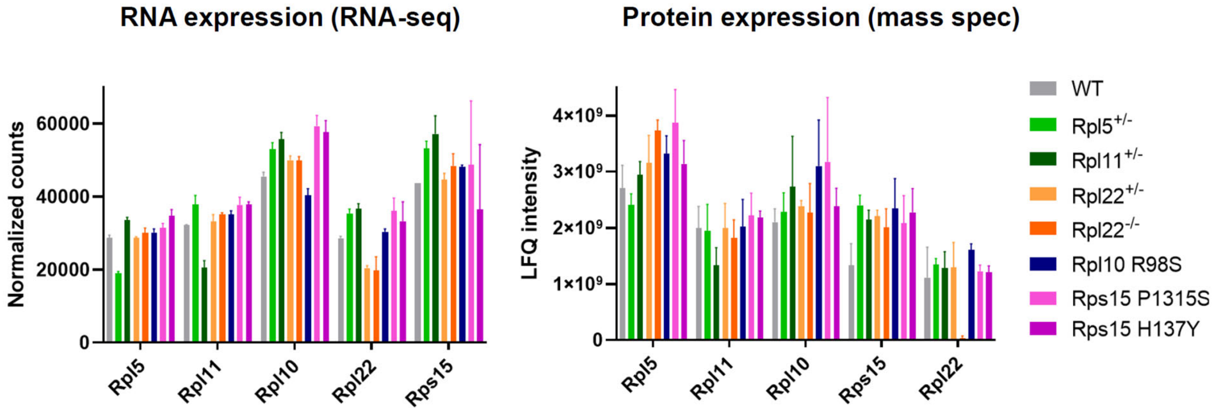

**Supplementary Figure 1. mRNA and protein levels of RP genes in RiboCancer cell line panel.** Left: mRNA levels of the indicated RP genes in the RiboCancer cell line panel as detected by RNA-seq. Right: protein levels of the indicated RP genes as detected by mass spectrometry proteomics analysis. The colors of the bars in the graph indicate the different genotypes modeled in the RiboCancer cell line panel as indicated in the legend on the right. Each bar in the plot represents the average value from three independent clones per genotype that were analyzed, with indication of the standard deviation (SD).

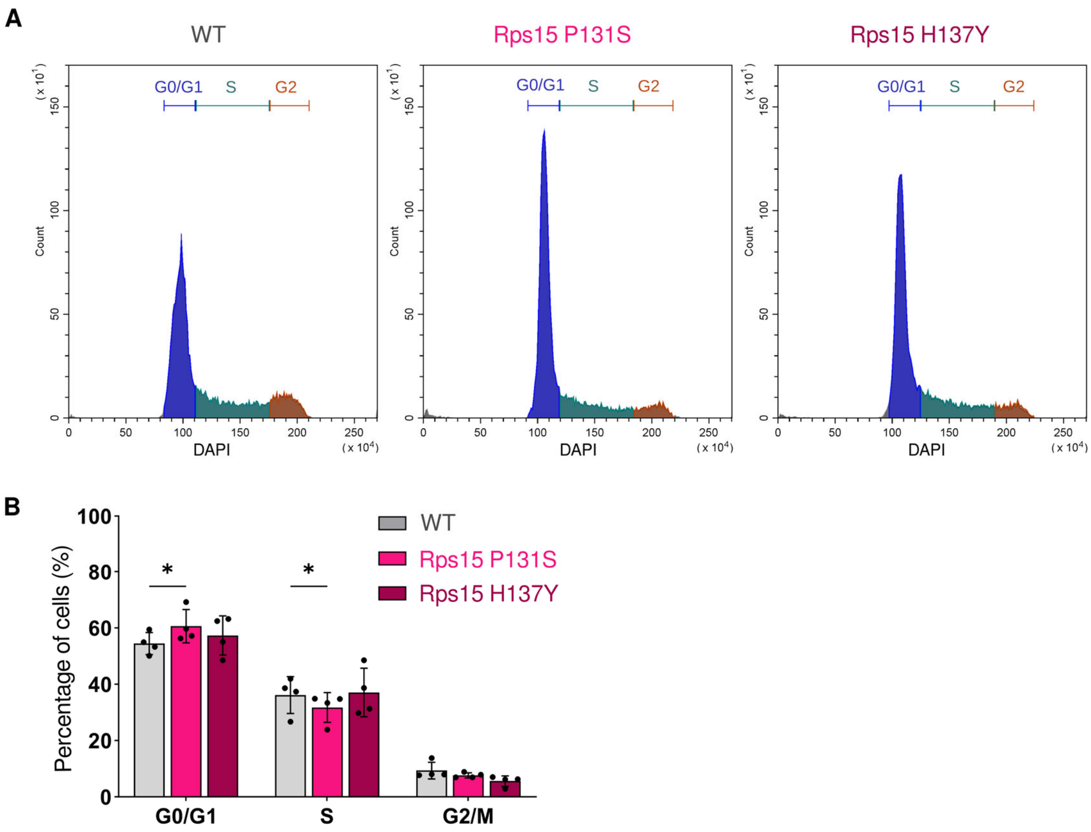

**Supplementary Figure 2. Rps15 point mutations lead to increased fraction of cells in G0/G1 phases.** **A)** DNA content analysis by flow cytometry of WT (left panel), Rps15 P131S (middle panel) and Rps15 H137Y (right panel) Ba/F3 cell lines. Signals corresponding to cell counts in different phases of the cycle are indicated: G0/G1 (blue), S (green), G2 (brown). **B)** Quantification of cell cycle distribution histograms (n=4 per genotype, three clones per genotype tested) shown in panel A. Data are presented as mean values +/− SD. Two-way ANOVA tests were used for statistics. * $p<0.05$.

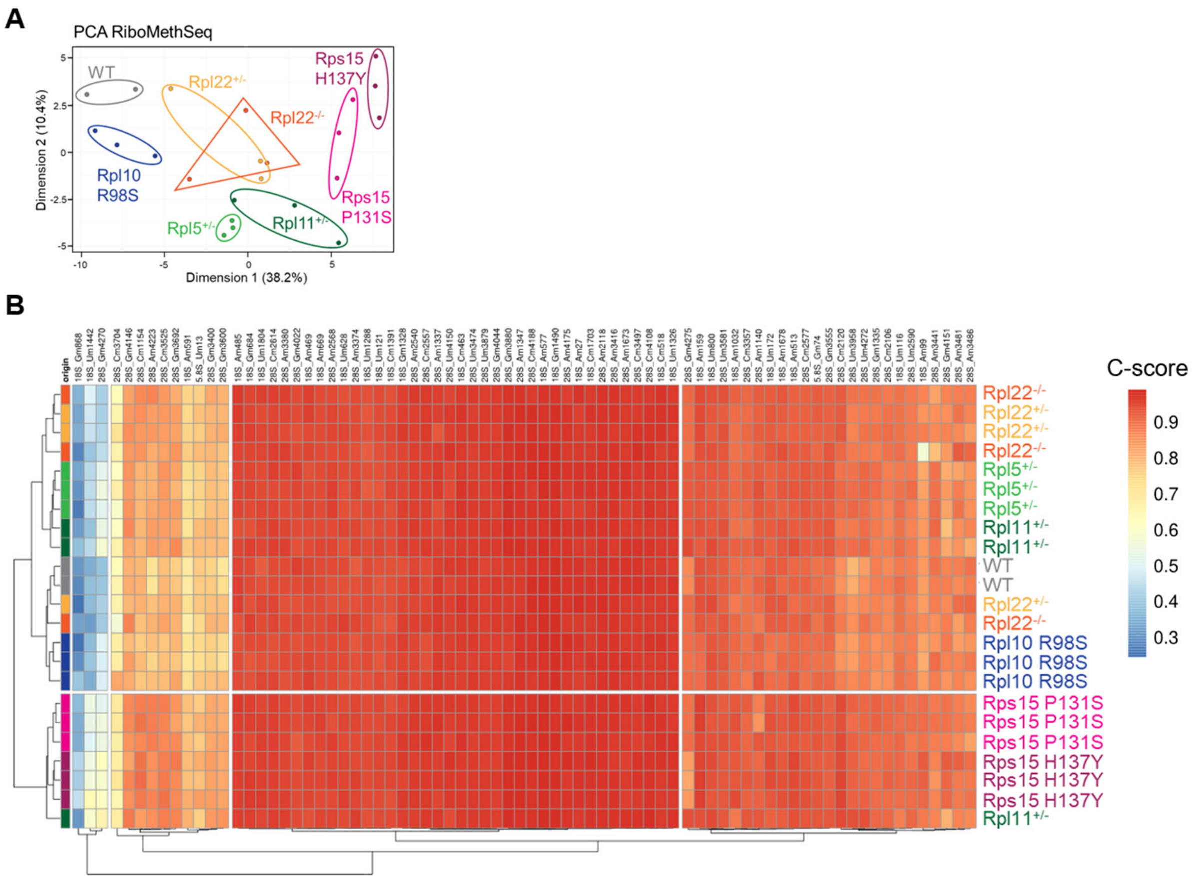

**Supplementary Figure 3. RiboMethSeq analysis of 2'-O-methylation of rRNA of ribosomes in the RiboCancer cell line panel. A)** PCA plot of RiboMethSeq data, each dot in the plot represents an independent single cell derived clone of the indicated genotype. **B)** Unsupervised hierarchical clustering of C-scores (representing the methylation level) at 76 known rRNA 2'O-ribose methylated sites. The genotypes of the cellular clones are indicated on the right. C-scores are represented by a color scale from 0 (dark blue) to 1 (red).

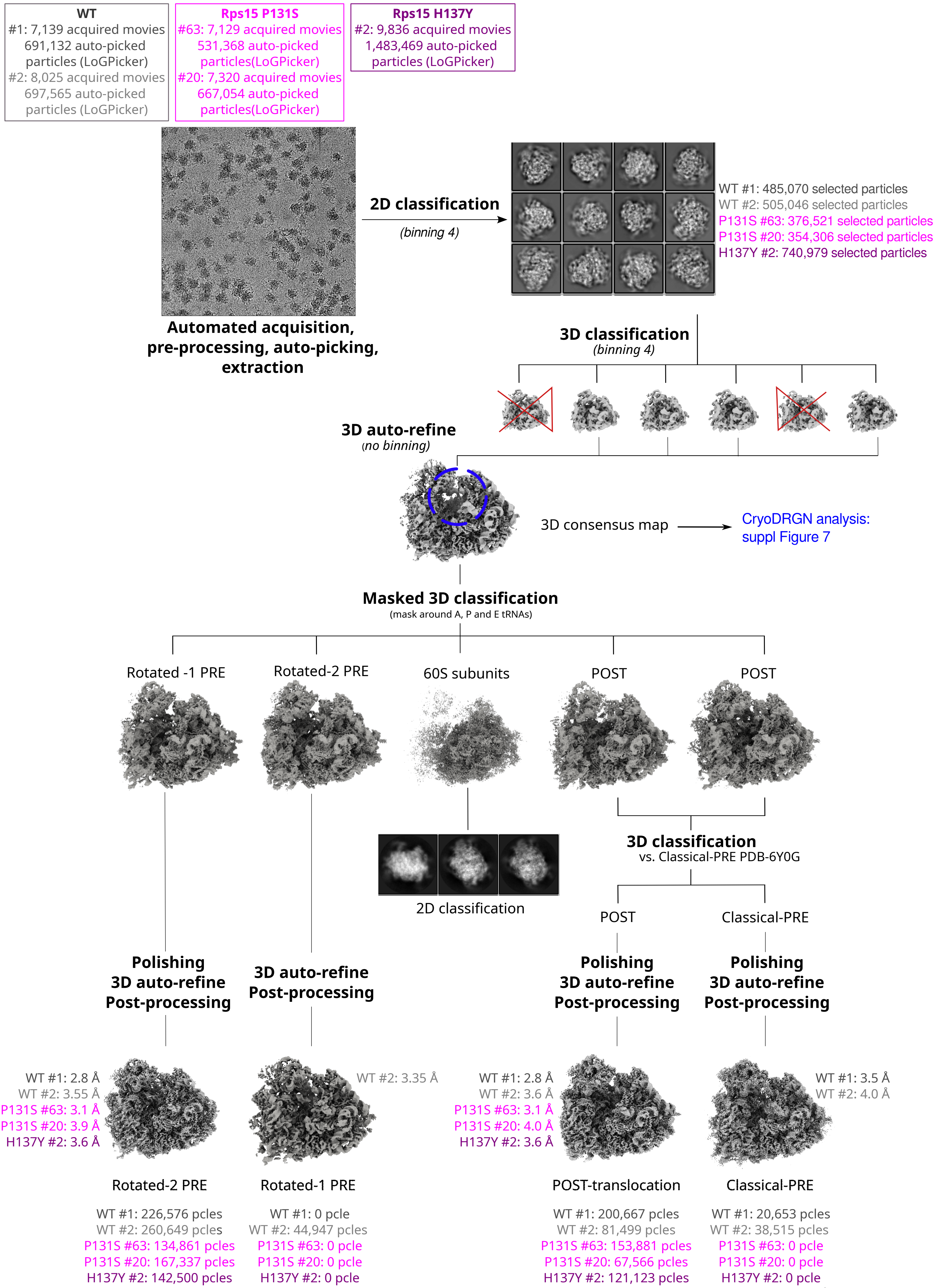

WT
#1: 7,139 acquired movies
691,132 auto-picked particles (LoGPicker)
#2: 8,025 acquired movies
697,565 auto-picked particles (LoGPicker)
Rps15 P131S
#63: 7,129 acquired movies
531,368 auto-picked particles(LoGPicker)
#20: 7,320 acquired movies
667,054 auto-picked particles(LoGPicker)
Rps15 H137Y
#2: 9,836 acquired movies
1,483,469 auto-picked particles (LoGPicker)
Automated acquisition, pre-processing, auto-picking, extraction
2D classification
(binning 4)
WT #1: 485,070 selected particles
WT #2: 505,046 selected particles
P131S #63: 376,521 selected particles
P131S #20: 354,306 selected particles
H137Y #2: 740,979 selected particles
3D classification
(binning 4)
3D auto-refine
(no binning)
3D consensus map
CryoDRGN analysis: suppl Figure 7
Masked 3D classification
(mask around A, P and E tRNAs)
Rotated -1 PRE
Rotated-2 PRE
60S subunits
POST
POST
2D classification
3D classification
vs. Classical-PRE PDB-6Y0G
POST
Classical-PRE
Polishing
3D auto-refine
Post-processing
3D auto-refine
Post-processing
Polishing
3D auto-refine
Post-processing
Polishing
3D auto-refine
Post-processing
WT #1: 2.8 Å
WT #2: 3.55 Å
P131S #63: 3.1 Å
P131S #20: 3.9 Å
H137Y #2: 3.6 Å
WT #2: 3.35 Å
WT #1: 2.8 Å
WT #2: 3.6 Å
P131S #63: 3.1 Å
P131S #20: 4.0 Å
H137Y #2: 3.6 Å
WT #1: 3.5 Å
WT #2: 4.0 Å
Rotated-2 PRE
Rotated-1 PRE
POST-translocation
Classical-PRE
WT #1: 226,576 pcles
WT #2: 260,649 pcles
P131S #63: 134,861 pcles
P131S #20: 167,337 pcles
H137Y #2: 142,500 pcles
WT #1: 0 pcle
WT #2: 44,947 pcles
P131S #63: 0 pcle
P131S #20: 0 pcle
H137Y #2: 0 pcle
WT #1: 200,667 pcles
WT #2: 81,499 pcles
P131S #63: 153,881 pcles
P131S #20: 67,566 pcles
H137Y #2: 121,123 pcles
WT #1: 20,653 pcles
WT #2: 38,515 pcles
P131S #63: 0 pcle
P131S #20: 0 pcle
H137Y #2: 0 pcle

**Supplementary Figure 4. Cryo-EM image processing pipeline, part 1**. SPA strategy applied to sort out structural states of the elongating ribosomes purified from WT (2 biological replicates using clones #1 and #2), Rps15 P131S (two biological replicates: clones #63 and #20) and H137Y (one experiment performed on clone #2) Ba/F3 cells. The number of initial images, extracted particles, final number of particles and resolutions for each type of ribosome are indicated in grey (WT), magenta (Rps15 P131S) and purple (H137Y).

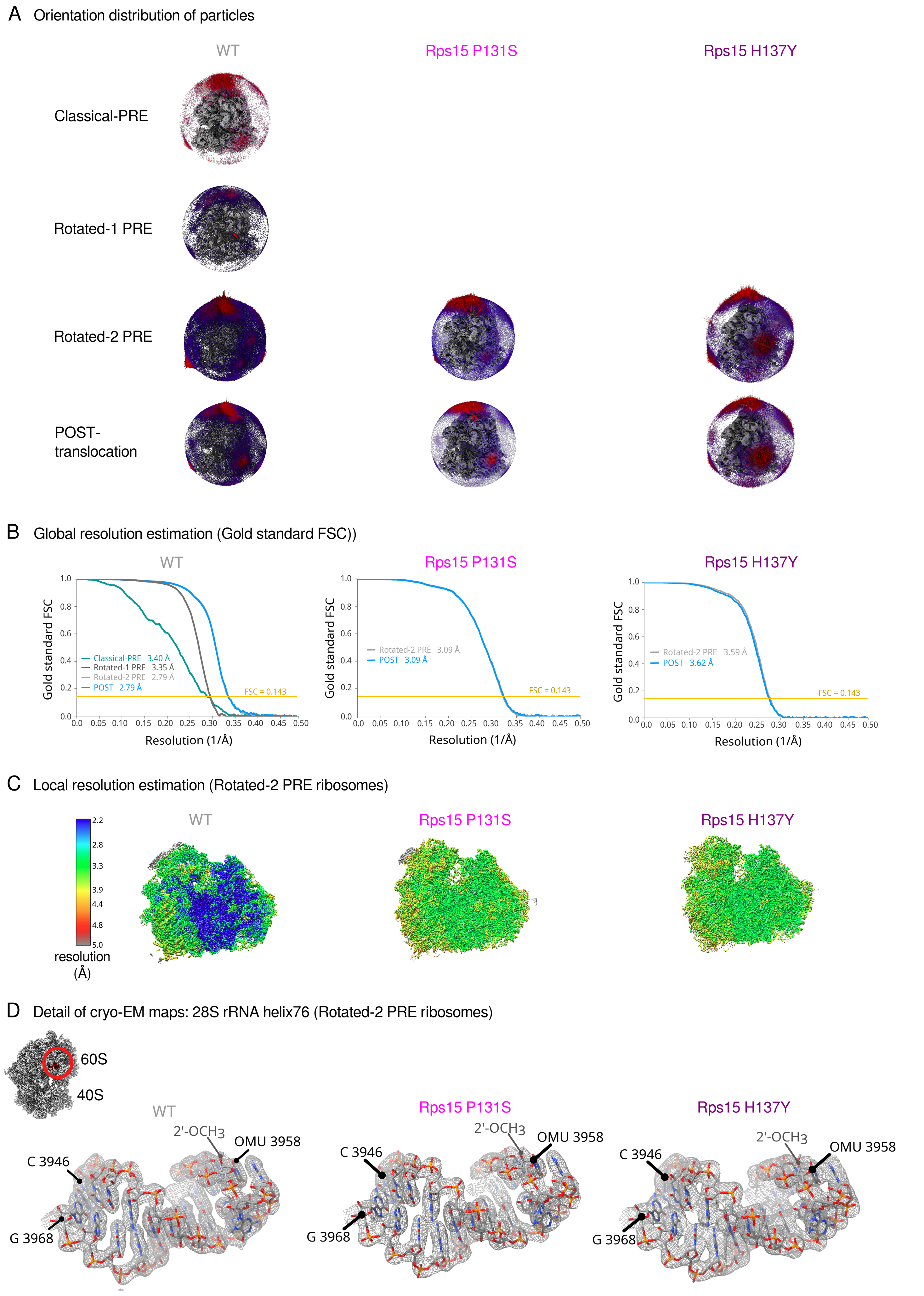
A Orientation distribution of particles
WT
Rps15 P131S
Rps15 H137Y
Classical-PRE
Rotated-1 PRE
Rotated-2 PRE
POST-
translocation
B Global resolution estimation (Gold standard FSC))
WT
Rps15 P131S
Rps15 H137Y
Gold standard FSC
Classical-PRE 3.40 Å
Rotated-1 PRE 3.35 Å
Rotated-2 PRE 2.79 Å
POST 2.79 Å
Rotated-2 PRE 3.09 Å
POST 3.09 Å
Rotated-2 PRE 3.59 Å
POST 3.62 Å
FSC = 0.143
Resolution (1/Å)
C Local resolution estimation (Rotated-2 PRE ribosomes)
WT
Rps15 P131S
Rps15 H137Y
2.2
2.8
3.3
3.9
4.4
4.8
5.0
resolution
(Å)
D Detail of cryo-EM maps: 28S rRNA helix76 (Rotated-2 PRE ribosomes)
60S
40S
WT
Rps15 P131S
Rps15 H137Y
2'-OCH3
OMU 3958
C 3946
G 3968

**Supplementary Figure 5. Cryo-EM image processing pipeline, part 2. A)** Euler angle distribution of the particles in each cryo-EM 3D map. **B)** Global resolution estimation (gold standard FSC) of each cryo-EM 3D map. **C)** Local resolution estimation of the cryo-EM 3D maps of Rotated-2 PRE ribosomes purified from Rps15 WT (left panel), Rps15 P131S (middle panel), Rps15 H137Y (right panel) Ba/F3 cells. **D)** Representative details of the cryo-EM 3D maps (grey mesh) and derived atomic model (colored sticks) in the vicinity of 28S rRNA helix 76. Only details of Rotated-2 PRE ribosomes purified from Rps15 WT (left panel), Rps15 P131S (middle panel), Rps15 H137Y (right panel) Ba/F3 cells are displayed. The 2'O-methylation (2'-OCH3 of Uridine 3958 (OMU 3958) is shown by arrows. C 3946 and G3968, at the other end of the helix, are also indicated.

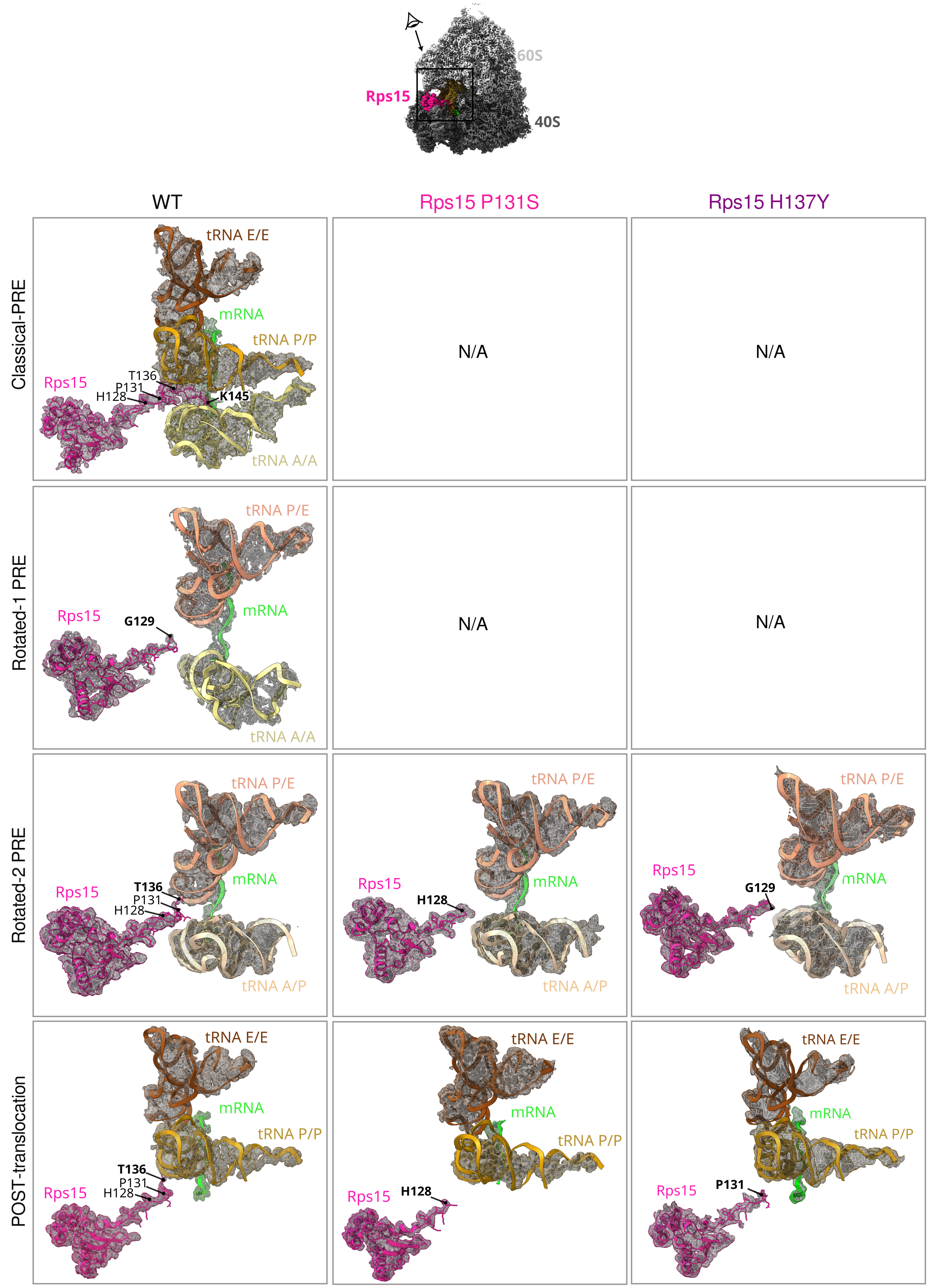
60S
40S
Rps15
WT
Rps15 P131S
Rps15 H137Y
Classical-PRE
tRNA E/E
mRNA
tRNA P/P
Rps15
T136
P131
H128
K145
tRNA A/A
N/A
N/A
Rotated-1 PRE
tRNA P/E
mRNA
Rps15
G129
tRNA A/A
N/A
N/A
Rotated-2 PRE
tRNA P/E
mRNA
Rps15
T136
P131
H128
tRNA A/P
tRNA P/E
Rps15
H128
mRNA
tRNA A/P
tRNA P/E
Rps15
G129
mRNA
tRNA A/P
POST-translocation
tRNA E/E
mRNA
tRNA P/P
T136
P131
H128
Rps15
tRNA E/E
mRNA
tRNA P/P
Rps15
H128
tRNA E/E
mRNA
tRNA P/P
Rps15
P131

**Supplementary Figure 6. Cryo-EM structures of Rps15 WT, P131S and H137Y ribosomes: zoom into the Rps15 region.** The elongation states (Classical-Pre, Rotated-1 or -2 PRE, and POST-translocation) for which we obtained near-atomic resolution 3D reconstructions are indicated on the left. Structural states that were not found in the Rps15-mutant datasets are indicated by N/A. As in Figure 3, Cryo-EM densities are displayed in transparent mesh, while modeled atomic structures are shown with ribbons and sticks. Rps15 is shown in magenta, mRNA in green, tRNA in yellow, golden and brown.

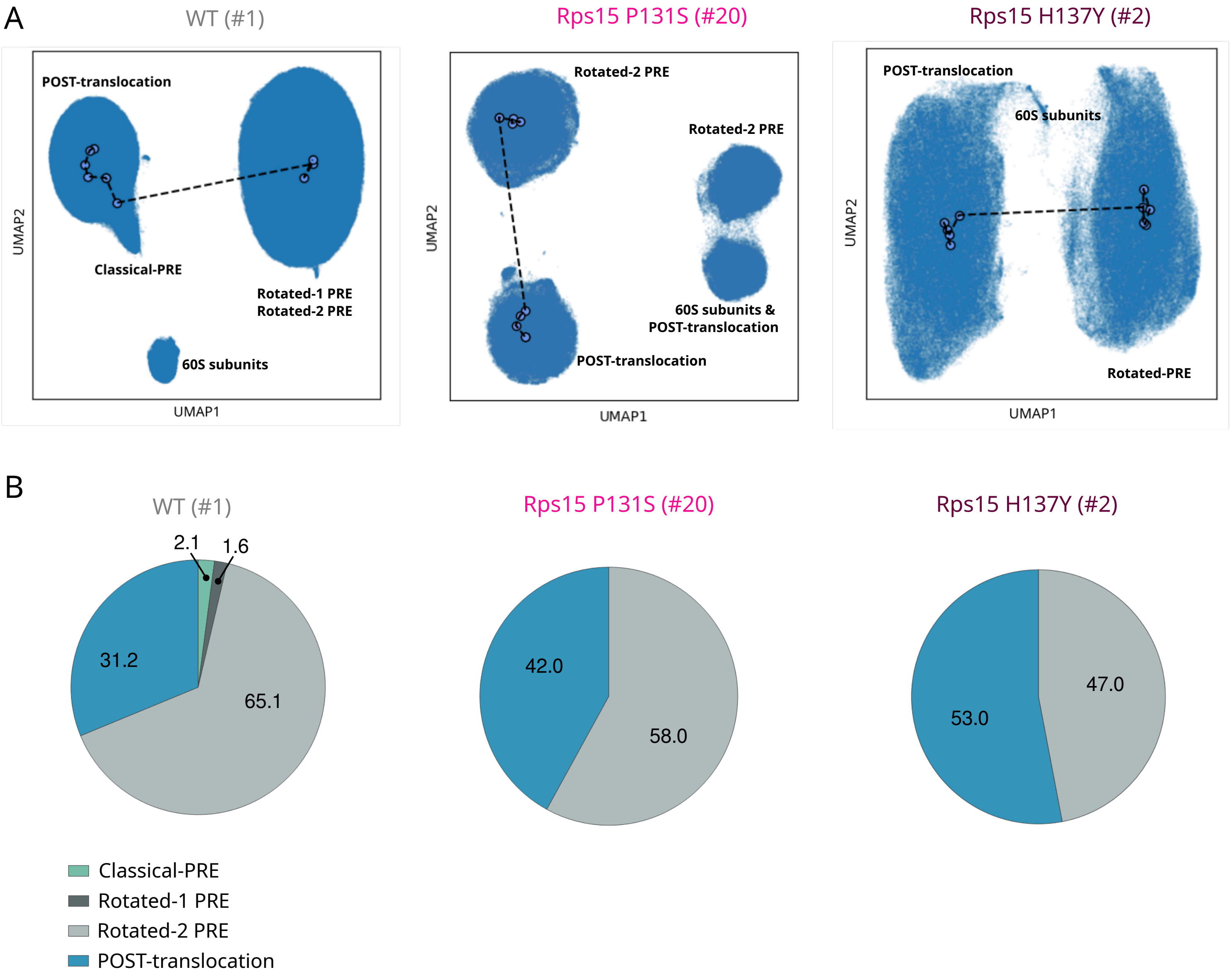

**Supplementary Figure 7. CryoDRGN analysis of the structural heterogeneity within Rps15 WT and Rps15 mutant cryo-EM datasets of translating ribosomes. A)** UMAP visualization of the latent space representation of particle images of WT (left panel), Rps15 P131S (middle panel) and Rps15 H137Y (left panel) translating ribosomes after training an 8-D latent variable model with cryoDRGN. Representative structural states (elongating ribosomes or free 60S subunits) are indicated next to the particles groups. **B)** For each cryo-EM dataset of ribosomes (WT: left panel, Rps15 P131S : middle and Rps15 H137Y: left panel), cryo-EM maps were reconstructed and grouped with cryoDRGN according to the elongation state, and the number of particles populating the Classical-PRE (green), rotated-1 PRE (dark grey), rotated-2 PRE (light grey) or POST-translocation (blue) states were counted, and represented as percentage of translating ribosomes. 60S subunits were excluded from count.

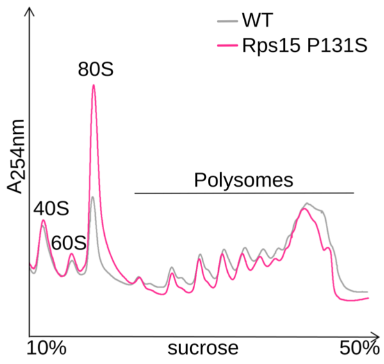

**Supplementary Figure 8. Polysome profiles from cytosolic extracts of WT and Rps15 P131S mutant cells.** These profiles were used to normalize the collision experiments presented in Figure 5B to the number of translating ribosomes (within polysomes) for each condition.

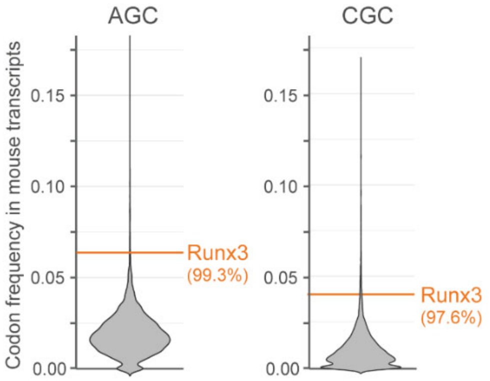

**Supplementary Figure 9. Runx3 is enriched for AGC and CGC codons.** Violin plot showing the frequencies of the AGC and CGC codons in mouse transcripts. The orange horizontal bar indicates the codon frequency in Runx3. The numbers between brackets indicate the percentile position of Runx3 for AGC and CGC codon frequency in the mouse transcriptome.

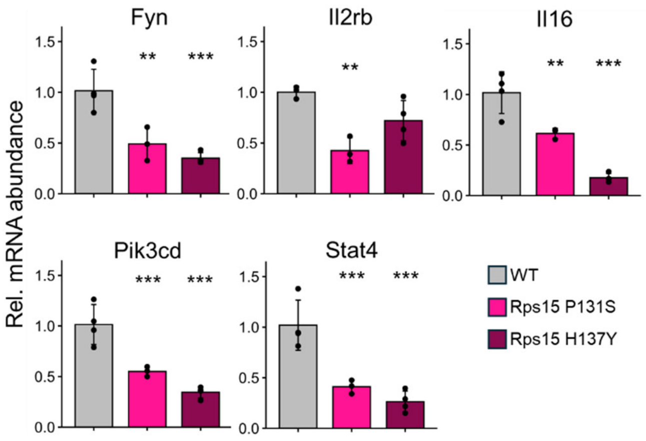

**Supplementary Figure 10. qPCR validation of Runx3 target genes mRNA expression in WT and Rps15 mutants.** Bar plot of relative mRNA abundance of selected Runx3 target genes when comparing Rps15 mutant Ba/F3 and WT cell lines. Data are presented as mean +/- SD from 4 WT, 3 Rps15 P131S and 4 Rps15 H137Y clones. Two-way ANOVA tests followed by Tuckey HSD post hoc test were used for statistics. Significance levels comparing mutants and WT cells are indicated above the bar plots.

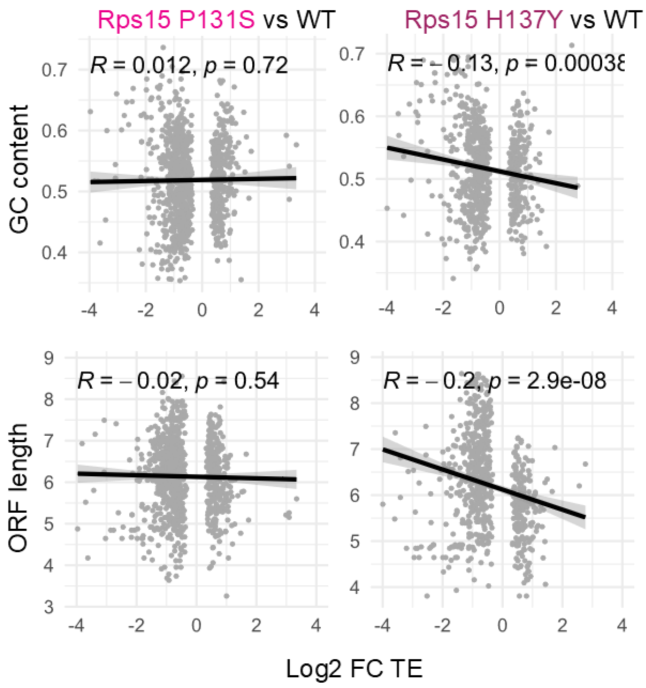

**Supplementary Figure 11. Correlation between translation efficiency (TE) and GC content, and between TE and ORF length.** Correlation analysis between either transcript GC content or ORF length, and translational efficiency (TE) fold change when comparing data obtained from Rps15 mutant and WT cells. Each dot in the plot represents a gene and shows the average value obtained from three independent clones per genotype that were analyzed. Only genes that were differentially translated were included in this analysis.

#### Supplementary tables

| Genotype | # generated single cell clones | Clone IDs |
|---|---|---|
| WT | 4 | **#1**; **#2**; **#3**; #A14 |
| $Rpl5^{+/-}$ | 4 | **#68; #76; #80;** #84 |
| Rpl10 R98S | 4 | #A22; **#A29; #A30; #A34** |
| $Rpl11^{+/-}$ | 4 | #21; **#22; #406; #414** |
| Rps15 P131S | 5 | **#20; #24;** #26; #60; **#63** |
| Rps15 H137Y | 4 | **#2; #6;** #15; **#16** |
| $Rpl22^{+/-}$ | 3 | **#13; #16; #60** |
| $Rpl22^{-/-}$ | 4 | **#30;** #604; **#649; #776** |

**Table S1. Overview of CRISPR-Cas9 engineered single cell clones and their IDs.** All clones in the table were used for the proteomics analysis. Cell clone IDs in bold were used for RNA-Seq and Ribo-Seq experiments.

**Table S2. (See separate file Supplementary_tables.xlsx) GSEA results of the mouse transcriptome after ranking all transcripts by their weighted 11-codon score.** The table reports results with a significant positive enrichment (pAdj < 0.05). Column L in the table reports the leading edge genes, illustrating a strong presence of histone transcripts in the gene sets with the strongest positive enrichment.

**Table S3. (See separate file Supplementary_tables.xlsx) GSEA results of translationally downregulated genes in Rps15 mutants.** The table reports the gene sets that are having a significant negative enrichment (pAdj < 0.05) for both analyzed Rps15 mutants (common enriched gene sets for Rps15 P131S vs WT and Rps15 H137Y vs WT). Negative enrichment indicates a reduction in TE in the Rps15 mutants as compared to WT. Columns F and I in the table report the leading edge genes, illustrating a strong presence of histone transcripts in the significant gene sets with the strongest reduction in TE.

**Table S4. (See separate file Supplementary_tables.xlsx) GSEA results of downregulated proteins in Rps15 mutants.** Results includes the terms that are commonly negatively enriched with a significant p value (pVal < 0.05) for both analyzed Rps15 mutants (Rps15 P131S vs WT and Rps15 H137Y vs WT). Negative enrichment indicates a reduction in protein levels in the Rps15 mutants as compared to WT.

| RP mutation | sgRNA sequence | ssODN sequence |
|---|---|---|
| Rpl5$^{+/-}$ | 5’-AATACTATCTATTACCTATG-3’ | None |
| Rpl10 R98S | 5’-TGATACGGATGACATGGAAA-3’ | 5’C*C*AACAAATACATGGTAAAGAGTTGTGGCAAGGATGGCTTTCATATCC GAGTGAGGCTCCATCCTTTTCATGTAATCAGTATCAACAAGATGTTGTCCTG TGCTGGGGCTGACAG*G*T 3’ |
| Rpl11$^{+/-}$ | 5’-CTCACCTTTGGAGAACACCG-3’ | None |
| Rpl22$^{+/-}$ or Rpl22$^{-/-}$ | 5’-ATACTCCTAGAAAAAGCTTG-3’ | None |
| Rps15 P131S | 5’-AAGCACGGCCGGCCCGGGAT-3’ | 5’A*A*AAACCCAGAATCCAGGGACCAAAACCAGTCTTTATTGGCCTCGGCT ACTTGAGGGGGATGAATCGGGAGGAGTGGGTGGCACCTATGCCGGACCG GCCGTGTTTCACGGGTTTGTAGGTGATGGAG*A*A 3’ |
| Rps15 H137Y | 5’-AAGCACGGCCGGCCCGGGAT-3’ | 5’A*A*AAACCCAGAATCCAGGGACCAAAACCAGTCTTTATTGGCCTCGGCT ACTTGAGGGGGATGAATCGGGAGGAGTAGGTGGCACCGATGCCGGGACG GCCGTGTTTCACGGGTTTGTAGGTGATGGAG*A*A 3’ |

**Table S5. Overview of sgRNA sequences and single-stranded oligodeoxynucleotide donor (ssODN) sequences used for CRISPR-Cas9 engineering.** For the ssODN sequences, synonymous and non-synonymous mutations are indicated in blue and red respectively, asterisks indicate phosphorothioate bonds.

| Gene | Forward primer | Reverse primer |
|---|---|---|
| Rpl5 | ACTCCAGACATGATGGAGGAGAT | TTGCCTTCTTTTGAGCAACCC |
| Rpl10 | TATATAAGGCGTGAGCAGGCG | CAATACCGGTAACACCGGGC |
| Rpl11 | TCCAGCAGAAGTACGATGGAATC | CACGCGCCTTTCACAAACAC |
| Rpl22 | CCTCGTGGTTAGCGGTCTTA | AAAGACCACCACATGGTCCG |
| Rps15 | TCCTTTTCCGAGTAACCGCC | AGCTGCATCAGTTGCTCATAGG |
| Pik3cd | CTCATTGGCAAAGGTCTCC | AAACTGGCGCATCTTAGTG |
| Stat4 | CTATCCAGGGACCTCTGGA | GCAGACACTTTGTGTTCCA |
| Il16 | TCTCCAAAGTCACCTTGCA | CAGCTCTGAAAGGTTGAGAG |
| Il2rb | CAACGAAACAATACCGGGAC | CTCCTTCATGGGATCTGCT |
| Fyn | GGACTAATTTGCAAGATTGCTG | GGAAACTTCGCACCTTGTC |
| Actb | CATTGCTGACAGGATGCAGAAGG | TGCTGGAAGGTGGACAGTGAGG |
| Hprt | CATTATGCCGAGGATTTGG | GCAAGTCTTTCAGTCCTGT |

**Table S6. Primer sequences (5'-3' direction) used for qRT-PCR.**

| Protein | Company | Catalogue number |
|---|---|---|
| RPL5 | Abcam | 157099 |
| RPL10 | Bio-Rad | VMA00429 |
| RPL11 | Abcam | ab79352 |
| RPS15 | Abcam | ab157193 |
| RPL22 | Santa Cruz | sc-373993 |
| RUNX3 | ThermoFischer Scientific | MA5-17169 |
| Histone H2B | Cell Signaling Technology | 12364T |
| Histone H3 | Abcam | ab1791 |
| Histone H4 | Cell Signaling Technology | 2592S |
| Vinculin | Sigma | V9131 |

**Table S7. Antibodies used for immunodetection.**

| | | | | | | | | | | | | |
|---|---|---|---|---|---|---|---|---|---|---|---|---|
| Microscope model | FEI Talos Arctica cryo-transmission electron microscope | | | | | | | | | | | |
| Detector model | Gatan K2 quantum summit direct electron detector | | | | | | | | | | | |
| Voltage (kV) | 200 | | | | | | | | | | | |
| | WT | | | | Rps15-P131S | | | | Rps15-H137Y | | | |
| Number of datasets | 2 | | | | 2 | | | | 1 | | | |
| Number of micrographs collected | 7,139<br>8,025 | | | | 7,129<br>7,320 | | | | 9,836 | | | |
| Number of selected particles | 691,132<br>697,427 | | | | 531,368<br>667,021 | | | | 1,483,469 | | | |
| Pixel size (Å) | 1.01 | | | | | | | | | | | |
| Defocus range (µm) | 0.8 - 2.8 | | | | | | | | | | | |
| Electron dose ($e^{-}Å^{-2}$) | 41.6<br>45 | | | | 35.2<br>45 | | | | 35.5 | | | |
| | CP | RP-1 | RP-2 | POST | CP | RP-1 | RP-2 | POST | CP | RP -1 | RP-2 | POST |
| EMDB entry of map | EMD-53473 | EMD-53333 | EMD-53310 | EMD-53262 | | | EMD-53427 | EMD-53307 | | | | |
| PDB entry of the full model | 9QZP | 9QSA | 9QQP | 9QOH | | | 9QWT | 9QQL | | | | |
| Final number of particles | 20,653<br>38,515 | 44,947 | 226,576<br>260,649 | 200,667<br>81,499 | | | 134,861<br>167,337 | 153,881<br>67,566 | | | 142,500 | 121,123 |
| Resolution (Å) (FSC threshold = 0.143) | 3.4<br>4 | 3.35 | 2.8<br>3.55 | 2.8<br>3.6 | | | 3.1<br>3.9 | 3.1<br>4 | | | 3.6 | 3.6 |
| Map sharpening B-factor (Å$^2$) | -61<br>-160 | -177 | -84<br>-183 | -83<br>-160 | | | -119<br>-198 | -112<br>-186 | | | -176 | -167 |

| Refinement and model validation statistics[a] | | | | | | | | |
|---|---|---|---|---|---|---|---|---|
| Model refinement resolution range (Å) | 15-3.4 | 15-3.3 | 15-2.8 | 15-2.8 | | | 15-3.1 | 15-3.1 |
| Model resolution (Å) (FSC threshold = 0.5) | 4.41 | 5.04 | 3.17 | 3.02 | | | 3.9 | 3.52 |
| Clashscore (all atoms) | 10.47 | 11.55 | 8.09 | 9.01 | | | 8.74 | 8.26 |
| MolProbity Score | 1.84 | 1.94 | 1.75 | 2.04 | | | 1.84 | 2.05 |
| Protein | | | | | | | | |
| Rmsd (bonds lengths, Å) | 0.002 | 0.004 | 0.003 | 0.004 | | | 0.007 | 0.004 |
| Rmsd (angles, °) | 0.582 | 0.614 | 0.582 | 0.621 | | | 0.674 | 0.616 |
| Ramachandran plot (%) | | | | | | | | |
| Favored | 100 | 99.9 | 99.9 | 100 | | | 100 | 100 |
| Outliers | 0 | 0.1 | 0.1 | 0 | | | 0 | 0 |
| RNA | | | | | | | | |
| Correct sugar puckers (%) | 100 | 100 | 100 | 100 | | | 100 | 100 |
| Good backbone conformation (%) | 78 | 78 | 78 | 78 | | | 80 | 79 |
| [a]Models were validated using MolProbity implemented in PHENIX.REFINE (Adams et al., 2010) | | | | | | | | |

**Table S8. Cryo-EM data collection, atomic models refinement and validation statistics**
